# Kinetochore Homeostasis is Maintained by Coordinated Chromatin Stabilization and Soluble Buffering

**DOI:** 10.64898/2026.01.07.698278

**Authors:** Shreyas Sridhar, Reiko Nakagawa, Tatsuo Fukagawa

**Affiliations:** Graduate School of Frontier Biosciences, The University of Osaka, Suita, Osaka, Japan; Laboratory for Cell-Free Protein Synthesis, RIKEN Center for Biosystems Dynamics Research (BDR), Kobe, Japan

**Keywords:** Kinetochore, Homeostasis, Chromosome segregation, Proximity labeling

## Abstract

Faithful chromosome segregation requires the coordinated assembly and maintenance of kinetochore complexes. However, the homeostatic mechanisms that maintain these multi-subunit assemblies remain unclear. CENP-T is a chromatin-bound linker that recruits the outer kinetochore modules via the CENP-T-W-S-X complex to form a functional kinetochore. Although CENP-T must be continuously replenished, how its steady-state levels are maintained is unknown. Here we demonstrate that CENP-T homeostasis is actively sustained by two spatially distinct pathways. Using covalent pulse-labeling, we find that CENP-T undergoes cell-cycle-coupled turnover rather than being stably inherited. *In situ* proximity labeling of chromatin-bound and soluble pools identifies regulators of CENP-T stability, including an unexpected role for the CENP-O complex in stabilizing soluble CENP-T. Mechanistically, CENP-S-X stabilizes chromatin-bound CENP-T through DNA-binding, whereas the CENP-O complex buffers a replenishment-competent CENP-T pool. Together, these conserved, spatially compartmentalized pathways establish an actively maintained homeostatic system that ensures robust kinetochore assembly and faithful chromosome segregation.

## INTRODUCTION

Cells rely on the precise homeostasis of multicomponent protein assemblies to execute essential functions, yet how such large molecular machines are dynamically maintained remains poorly understood. This challenge is particularly acute for the kinetochore, a macromolecular machine comprising over 100 proteins that connects centromeric chromatin with spindle microtubules to ensure faithful chromosome segregation.^1–3^ Errors in kinetochore assembly drive chromosome mis-segregation and aneuploidy,^4–9^ established hallmarks of tumorigenesis.^10,11^ Despite the structural composition of the kinetochore being increasingly well defined,^12–16^ the regulatory principles that preserve its stoichiometry and functional integrity across the cell cycle remain unresolved, reflecting broader principles of cellular homeostasis and quality control systems that remain incompletely understood.

At the foundation of kinetochore architecture in vertebrates lies the 16-member constitutive centromere-associated network (CCAN), comprised of CENP-C, CENP-H-I-K-M, CENP-L-N, CENP-T-W-S-X and CENP-O-P-Q-U-R subcomplexes.^3,17^ The CCAN links centromeric CENP-A nucleosomes ^18–23^ to the outer kinetochore KMN (KNL1, Mis12 and Ndc80 complexes) network, which directly binds to spindle microtubules.^3,23,24^ Within this network, CENP-T occupies a central position in forming a functional kinetochore as it acts both as a structural component capable of direct interaction with centromeric DNA and as the essential platform for recruiting the KMN network and in turn the spindle assembly checkpoint machinery.^25–33^ The CENP-T complex is comprised of two related, but functionally distinct subcomplexes - CENP-T-CENP-W and CENP-S-CENP-X^34^ - heterodimeric histone-fold pairs that assemble into a nucleosome-like tetramers capable of wrapping DNA.^25,35^ The CENP-T complex and the histone H3 variant CENP-A are the only kinetochore components that possess direct nucleosome-like DNA-binding properties.^25,35^

Recent cryo-EM studies have provided detailed snapshots of CCAN architecture.^12,36^ Although these studies provide useful structural information, they offer only limited insight into how the intricate CCAN architecture is established, renewed, and maintained after each cell division. CENP-A deposition is tightly regulated at the G1 phase in vertebrates,^37,38^ whereas the regulatory pathways governing CENP-T turnover and replenishment remain poorly defined leaving a fundamental gap in understanding kinetochore maintenance. This raises a central question of how CENP-T homeostasis is maintained to ensure robust kinetochore function each cell cycle.

Here, we address this challenge by integrating systematic proximity labeling, functional genomics, and quantitative microscopy approaches. We define the turnover dynamics of the CENP-T complex and identify the molecular factors responsible for the regulation of both its soluble and chromatin-bound pools. Simultaneous disruption of these pathways causes loss of CENP-T from the kinetochore and leads to severe mitotic defects, revealing that kinetochore assembly relies on sophisticated, evolutionarily conserved quality-control mechanisms that balance soluble and chromatin-bound protein pools. These findings reshape classical notions of constitutive kinetochore inheritance and uncover broader principles of spatially compartmentalized molecular homeostasis that ensure genome stability across cell divisions.

## RESULTS

### CENP-T complex undergoes rapid turnover and assembles during a defined S/G2 window

Despite the nucleosome-like CENP-T complex’s central role in kinetochore function, its dynamic behavior has remained poorly defined. In particular, it is unclear whether its two functionally distinct subcomplexes - the CENP-T-W and CENP-S-X histone-fold pairs - assemble or turn over independently or as a coordinated unit each cell cycle. To directly dissect these dynamics, we combined CRISPR-based genome editing with SNAP^39^ and Halo^40^ covalent labeling systems in chicken DT40 cells. All endogenous CENP-T alleles were tagged with mScarlet-3xHA-SNAP-CENP-T (mS-SNAP-CENP-T) to monitor the dynamics of the CENP-T-W subcomplex (Figures S1A and S1B). In parallel, CENP-S was endogenously tagged with CENP-S-2xMyc-Halo (CENP-S-Halo) to track the dynamics of the CENP-S-X subcomplex (Figure S1C). To assess inheritance and stability, we performed pulse-chase labeling and tracked signal retention across multiple cell divisions (Figure 1A). Unlike the epigenetic centromeric mark CENP-A, which is diluted but not replaced,^37^ both CENP-T and CENP-S underwent rapid cell-cycle-coupled turnover, losing ∼80% and ∼75% of pulse-labeled signal, respectively, with each cell cycle (Figures 1B and 1C). These measurements indicate that the CENP-T complex is not stably inherited but must be actively renewed.

**Figure 1.**
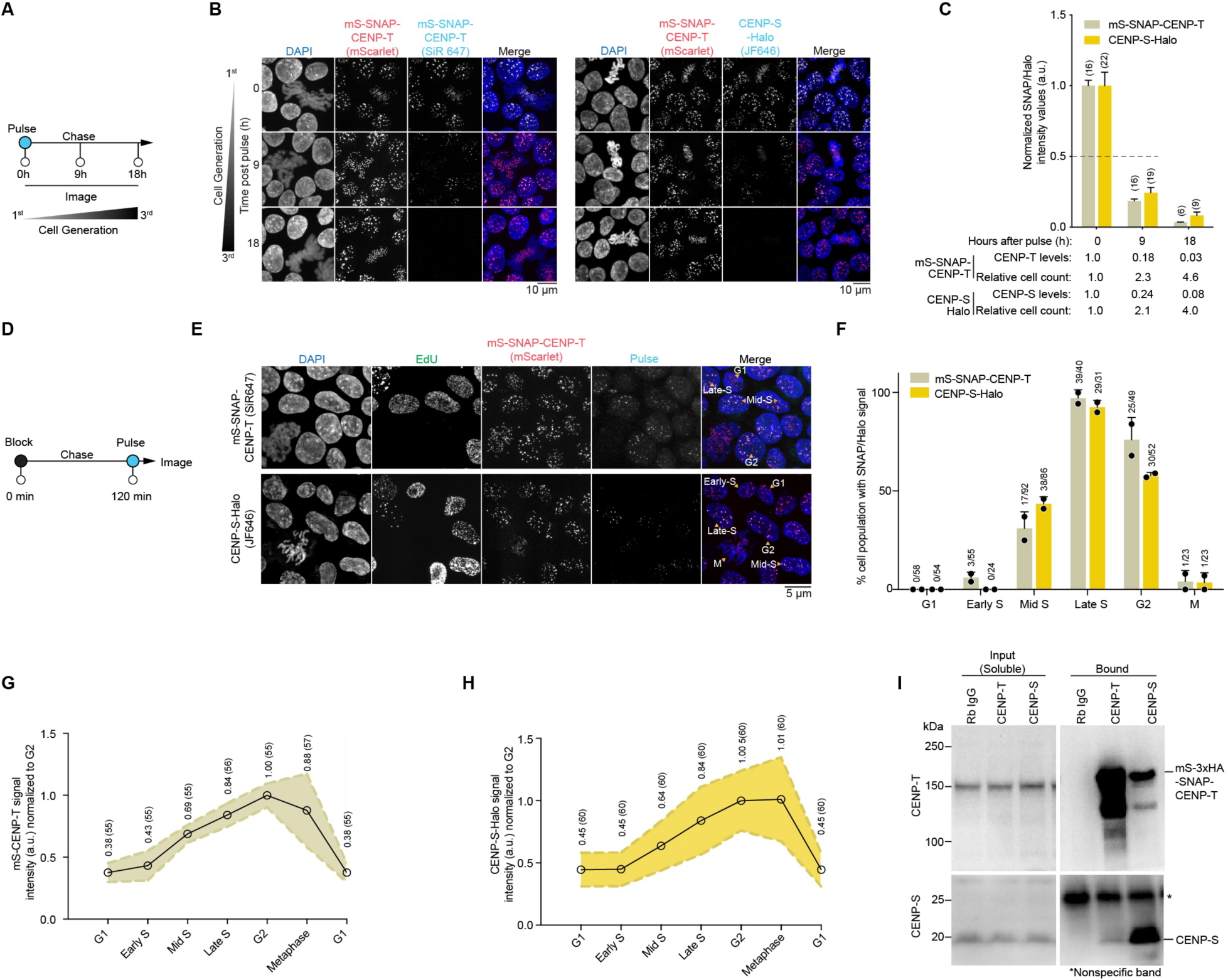
The CENP-T complex undergoes dynamic renewal each cell cycle during S-G2 stages. (A) Schematic of SNAP/Halo pulse chase assay. Total cellular mScarlet(mS)-SNAP-CENP-T or CENP-S-Halo pool was pulsed at 0 h with SNAP-Cell 647-SiR or JF646, respectively, and was subsequently washed. Labelled CENP-T complex signal was chased over 2 successive cell divisions and imaged, DT40 doubling time is ∼9 h. (B and C) Localization of SNAP or Halo labelled CENP-T or CENP-S, respectively, over 0, 9 and 18 h of the pulse chase assay. SNAP or Halo labelled signal intensities in mitotic cells were quantified in (C). Number (*n*) of mitotic cells quantified is labelled in parenthesis, mean ± standard error of mean (SEM). (D) Schematic of SNAP/Halo loading assay. Total cellular mS-SNAP-CENP-T or CENP-S-Halo pool was labelled with a non-fluorescent reagent at 0 h and subsequently washed. Cells were chased for 2 h to allow for new unlabeled protein synthesis and was pulsed with SNAP-Cell 647-SiR or JF646 for mS-SNAP-CENP-T or CENP-S-Halo, respectively, and imaged to visualize the localization of newly synthesized protein. (E and F) Localization of SNAP or Halo labelled CENP-T or CENP-S, respectively, following the 2 h chase. 5-ethynyl-2’-deoxyuridine (EdU) labeling was performed to track cell cycle stages as described in Figure S1D. The number of cells positive for SNAP/Halo signal across the cell cycle were quantified in (F). *n* = 2, ≥ 270 cells, mean ± standard deviation (SD). (G) mScarlet signal intensities of mS-SNAP-CENP-T, as represented in Figure S1E, was quantified across cell cycle stages. Number (*n*) of cells quantified is labelled in parenthesis, mean ± SD. (H) JF646 labeled CENP-S-Halo signal intensities, as represented in Figure S1E, was quantified across cell cycle stages. Mean intensity is reported and the number (*n*) of cells quantified is labelled in parenthesis, mean ± SD. (I) Western blot of CENP-T and CENP-S immunoprecipitation (IP) from the soluble pool. Proteins were extracted following cell fractionation and IP was performed using anti-CENP-T, anti-CENP-S or control rabbit IgG. *n* = 2. Cell fractionation efficiency is shown in Figure S1F.

To determine when new CENP-T and CENP-S molecules are assembled, we performed block-chase-pulse experiments in cells co-labeled with EdU to determine cell-cycle stage (Figures 1D and S1D). Newly incorporated CENP-T and CENP-S appeared almost exclusively during S and G2, with late-S and G2 cells showing the strongest signals (Figures 1E and 1F). This sharply contrasts with CENP-A, whose deposition is confined to G1.^37,38^ We did not observe detectable CENP-T turnover during mitosis, supporting stable retention on mitotic kinetochores, and consistent with a previous report.^41^ Quantification of total kinetochore levels revealed a parallel increase of CENP-T and CENP-S beginning at S-phase entry and peaking in G2 (Figures 1G, 1H and S1E). Thus, both CENP-T subcomplexes share a coordinated and temporally restricted S/G2 assembly program.

Given their similar turnover dynamics, we confirmed that CENP-T complex subunits can associate when co-localized. Tethering either CENP-W or CENP-S to an ectopic chromosomal array recruited endogenous CENP-T (Figures S1F-G). To test whether such interactions also occur under physiological conditions before chromatin incorporation, we examined soluble cellular fractions by co-immunoprecipitation (co-IP). This analysis revealed clear pre-assembly between CENP-T and CENP-S (Figures 1I and S1I), supporting the presence of a soluble CENP-T complex.

Together, these results indicate that the CENP-T complex is dynamically renewed and that its subunits can associate in the soluble fraction before their incorporation at centromeres during a defined S/G2 window each cell cycle.

### CENP-S DNA-binding is essential for stable CENP-T maintenance at kinetochores

CENP-S was shown previously to be non-essential for cell viability, but its loss disrupts the localization of its binding partner CENP-X in chicken DT40 cells.^42^ In vitro, CENP-S-X remodels the DNA-binding behavior of CENP-T-W, converting its weak, irregular DNA contacts into a discrete, stoichiometric binding mode that emerges on DNA substrates >80 bp.^25^ Notably, whereas CENP-T-W or CENP-S-X alone introduce negative supercoils, the assembled CENP-T-W-S-X heterotetramer induces positive supercoiling, suggesting that subcomplex assembly remodels DNA into a distinct higher-order conformation.^35^ However, how CENP-S-X contributes to CENP-T dynamics and kinetochore integrity in vivo has remained unclear (Figure 2A).

**Figure 2.**
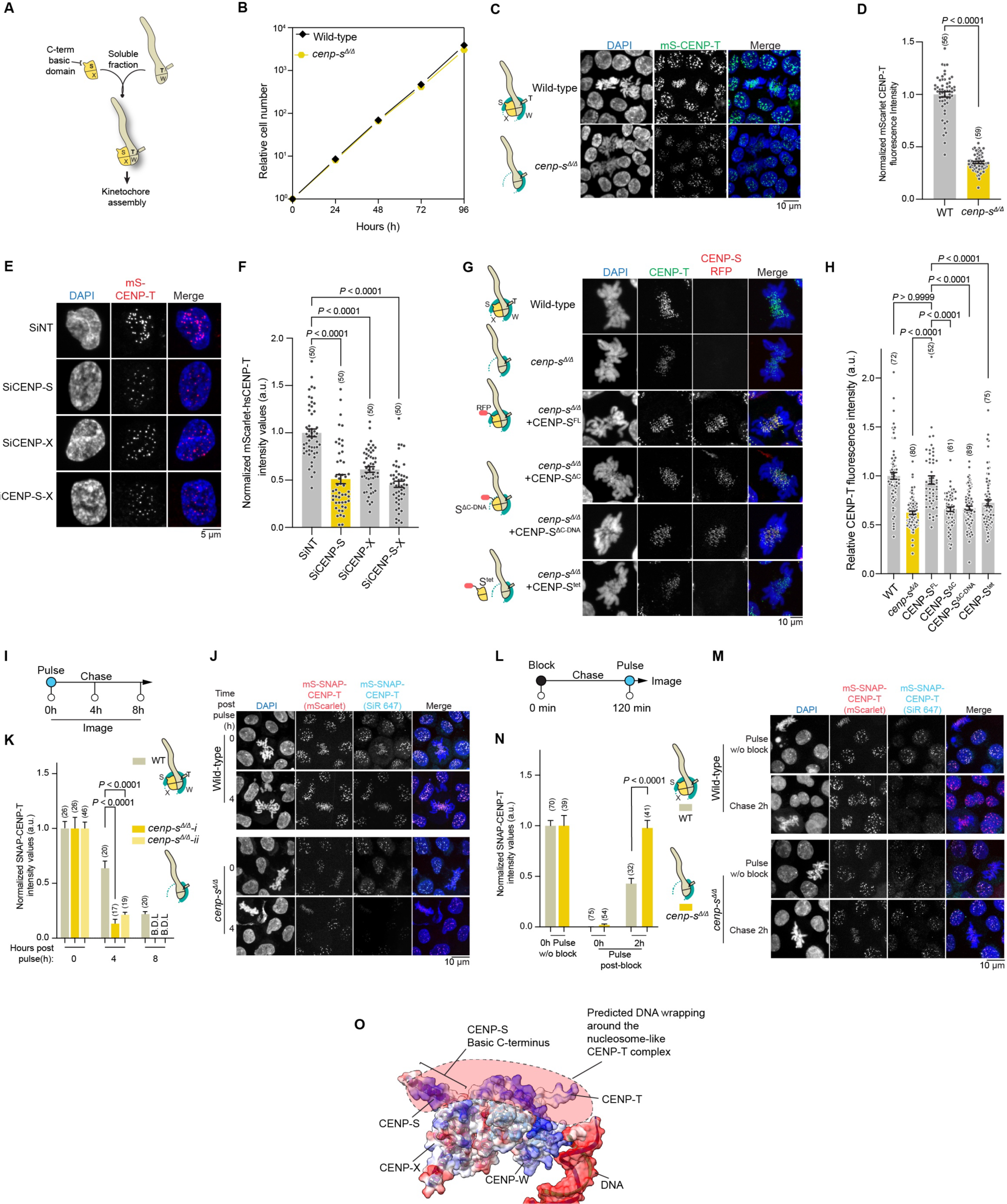
CENP-S promotes CENP-T complex stability at kinetochores through enhanced centromere DNA binding. (A) Schematic of CENP-T complex formation in the soluble pool, followed by its kinetochore assembly. (B) Growth of wild-type (CL18) and *cenp-s^1/1^* cells. The cell numbers were normalized to those at 0 h for each cell line. (C and D) Localization of CENP-T in wild-type (WT) and *cenp-s^1/1^* cells. mS-CENP-T intensities in mitotic cells were quantified in (D). Statistical analysis was performed with a non-parametric t-test comparing two unpaired groups (Mann-Whitney test), number (*n*) of mitotic cells quantified is labelled in parenthesis, mean ± SEM, *P* values are reported as indicated. (E and F) Localization of CENP-T in human RPE-1 cells following treatment of small interfering RNA (SiRNA) towards control (non-target guide, SiNT), CENP-S (SiCENP-S), CENP-X (siCENP-X) or both CENP-S and CENP-X (SiCENP-S-X). CENP-T intensity in interphase cells were quantified in (F). Statistical analysis was performed with Kruskal–Wallis one-way analysis followed by Dunn’s multiple comparison test, number (*n*) of interphase cells quantified is labelled in parenthesis, mean ± SEM, *P* values are reported as indicated. (G and H) Localization of CENP-T and CENP-S in WT (CL18), *cenp-s^1/1^ or cenp-s^1/1^* complimented with CENP-S^FL^, CENP-S^1C^, CENP-S^1C+DNA^ or CENP-S^tet^ mutant constructs. CENP-T signal in mitotic cells was quantified and internally normalized to co-mixed reference cells in (H). Statistical analysis was performed with Kruskal–Wallis one-way analysis followed by Dunn’s multiple comparison test, number (*n*) of mitotic cells quantified is labelled in parenthesis, mean ± SEM, *P* values are reported as indicated. (I) Schematic of SNAP pulse chase assay. Total cellular mS-SNAP-CENP-T was pulsed at 0 h with SNAP-Cell 647-SiR and was subsequently washed. SNAP Labelled CENP-T signal was chased over 8 h. (J and K) Localization of SNAP labeled CENP-T at 0, 4 and 8 h of the pulse chase assay in WT and *cenp-s^1/1^* cells. SNAP labelled signal intensities in mitotic cells were quantified in (K). Statistical analysis was performed with Kruskal–Wallis one-way analysis followed by Dunn’s multiple comparison test, number (*n*) of mitotic cells quantified is labelled in parenthesis, mean ± SEM, *P* values are reported as indicated. SNAP-labelled CENP-T in *cenp-s^1/1^* cells were below detectable levels (B.D.L) at 8 h. (L) Schematic of SNAP loading assay. Total cellular mS-SNAP-CENP-T was labelled with a non-fluorescent reagent at 0 h and subsequently washed. Cells were chased for 2 h to allow for new unlabeled protein synthesis and was pulsed with SNAP-Cell 647-SiR, imaged to visualize the localization of newly synthesized protein. (M and N) Localization of SNAP pulsed CENP-T prior to block at 0 h and 2 h post block in wild-type and *cenp-s^1/1^* cells. SNAP labeled CENP-T intensity in G2 were quantified and normalized to its respective 0 h pulse in (N). Statistical analysis was performed with a non-parametric t-test comparing two unpaired groups (Mann-Whitney test), number (*n*) of cells quantified is labelled in parenthesis, mean ± SEM, *P* values are reported as indicated. (O) Schematic model describing the potential wrapping of centromeric DNA around the CENP-T nucleosome-like particle enhanced by the basic C-terminal tail and other basic residues of CENP-S. The molecular surface is colored by electrostatic potential, with red and blue representing regions of negative and positive charge, respectively. CENP-T complex DNA binding structure was adapted from Yatskevich, S. et al.^13^

To test this, we generated a CENP-S DT40 knockout (*cenp-s^1/1^*, KO) cells in the mS-SNAP-CENP-T background (Figure S2A). Consistent with the prior work, CENP-S loss was tolerated and did not impair cell proliferation (Figure 2B).^42^ As CENP-T provides the core platform for outer kinetochore recruitment,^6,23^ its levels are generally thought to be tightly constrained.^29^ However, unexpectedly, endogenous mScarlet-CENP-T levels were markedly reduced (by ∼65%) in *cenp-s^1/1^* cells (Figures 2C and 2D). Using immunofluorescence (IF) with internal normalization, the decrease appeared more modest (∼35%) (Figure S2B), likely reflecting increased antibody accessibility in the absence of CENP-S.

To test for conservation in human cells, we depleted CENP-S and CENP-X by small interfering RNA (siRNA) in human retinal pigment epithelial (RPE-1) cells expressing mScarlet-CENP-T and GFP-mini auxin-inducible degron (mAID)-CENP-T.^29,43^ Acute degradation of GFP-mAID-CENP-T 24 h prior to analysis ensured uniform quantification of mScarlet-CENP-T (Figure S2C). As in DT40 cells, CENP-T levels were significantly reduced, supporting a conserved role for CENP-S in maintaining CENP-T at vertebrate kinetochores (Figures 2E, 2F and S2D).

We next analyzed whether CENP-S stabilizes CENP-T through DNA-binding or via tetramer formation with CENP-T-W. We expressed CENP-S DNA-binding mutants,^25^ including CENP-S^1C^ (lacking the C-terminal basic region, amino acids (aa) 107-139) or combining it with additional basic DNA-binding interface residues of R15A, K41A, K70A in CENP-S^1C-DNA^ and a tetramer-interface mutant, CENP-S^tet^ (F65E, H68E, L81R, R84E), which fails to localize to kinetochores due to impaired tetramerization with CENP-T-W, in the *cenp-s^1/1^* cells. Both DNA-binding mutants, CENP-S^1C^ and CENP-S^1C-DNA^, failed to restore normal CENP-T levels, phenocopying *cenp-s^1/1^* cells. The CENP-S^tet^ mutant behaved similarly to DNA-binding mutants, indicating that DNA-binding, not tetramer formation, is the primary determinant of CENP-T stabilization (Figures 2G and 2H).

Together, these results demonstrate that CENP-S preserves CENP-T abundance at the kinetochore primarily through its DNA-binding activity.

### CENP-S safeguards CENP-T through a post-assembly stabilization step

To determine whether the reduction in CENP-T levels upon CENP-S loss reflects impaired kinetochore maintenance, impaired kinetochore recruitment, or both, we measured CENP-T turnover in *cenp-s^1/1^* cells using a SNAP pulse-chase assay (Figure 2I). Labeled CENP-T decayed more rapidly in *cenp-s^1/1^* cells than wild type, revealing that CENP-S is essential for stable retention of CENP-T at kinetochores (Figures 2J and 2K). In contrast, CENP-T assembly remained robust in *cenp-s^1/1^* cells with newly synthesized CENP-T being incorporated efficiently into kinetochores (Figures 2L-2N). This demonstrates that CENP-S is dispensable for CENP-T incorporation but critical for post-assembly stability. Together, these findings identify a conserved stabilization step in CENP-T regulation that depends on CENP-S DNA-binding. By ensuring the continued retention of CENP-T after its incorporation, CENP-S provides a dynamic safeguard that complements and builds upon the DNA-interaction interface observed in recent CCAN cryo-EM structures^13,44^ to support kinetochore homeostasis throughout the cell cycle (Figure 2O).

### Proximity labeling reveals CENP-T interactome *in situ*

The active assembly of CENP-T following its increased turnover in *cenp-s^1/1^* cells is sufficient to maintain cell viability. This suggests that additional mechanisms help preserve functional CENP-T levels at kinetochores (Figure 3A). Structural studies identified the C-terminal α4-5 helices of the CENP-T histone fold, the histone-fold extension (HFE), as the interaction site for the CENP-H-I-K-M complex, critical for CENP-T localization.^13,45–47^ Additionally, mutations in the DNA-binding interface of CENP-W similarly disrupt CENP-T localization despite intact CENP-H-I-K-M binding,^45^ highlighting the combined importance of DNA engagement and protein interactions. Indeed, replacing the native histone-fold core (α1-3) with basic DNA-binding domains from Ki-67 ^48^ or the fungal linker Bridgin ^49^ failed to restore CENP-T localization or function (Figures S3A-S3E), demonstrating that the native CENP-T histone fold, or factors engaging with it, is indispensable for proper kinetochore activity.

**Figure 3.**
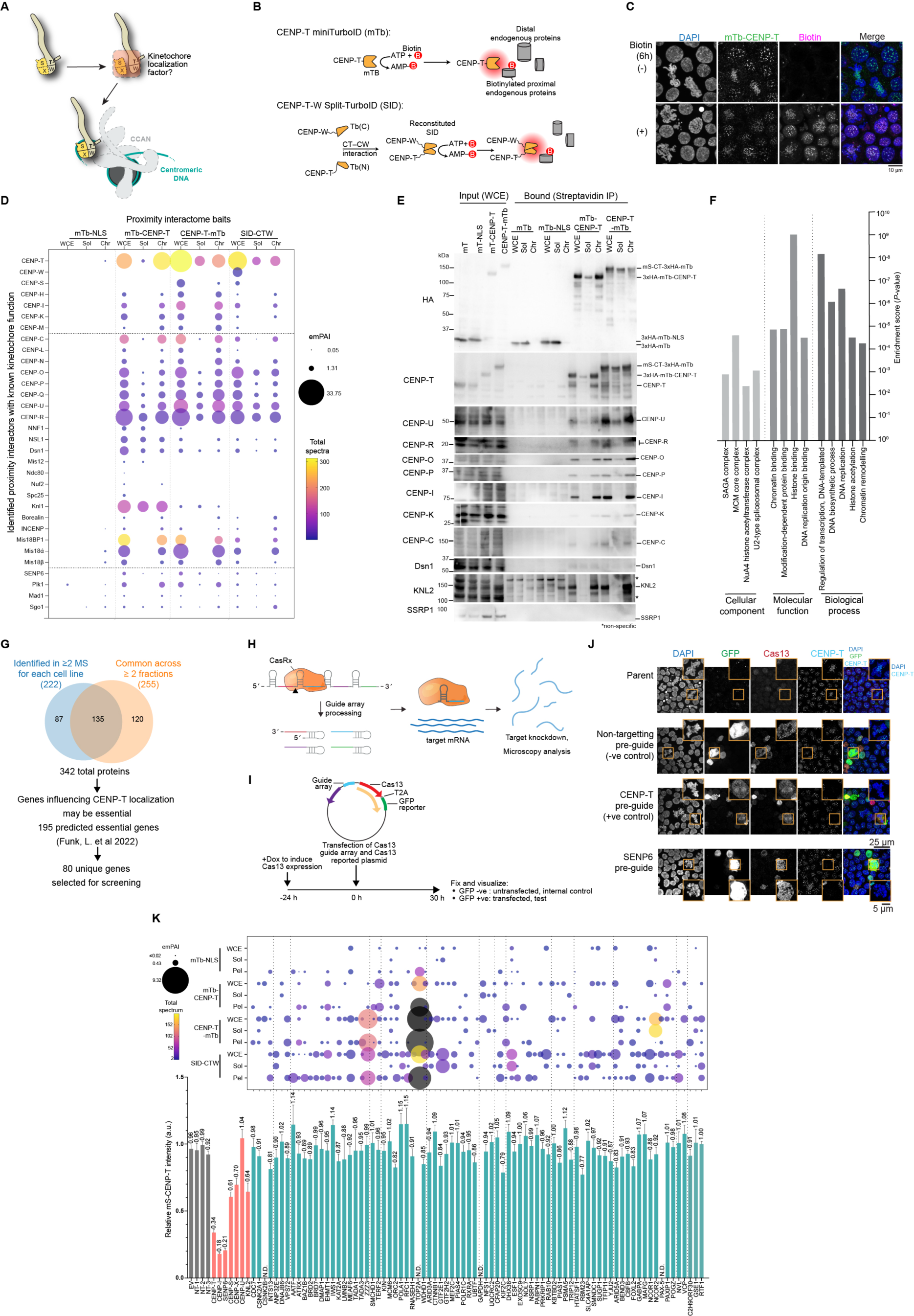
Mapping the CENP-T complex interaction landscape across kinetochore bound and unbound pools. (A) Schematic describing the likely presence of undescribed factors that aid in kinetochore localization of the CENP-T complex. (B) Schematic of TurboID (Tb)-based biotin proximity labeling using miniTurboID (mTb) and split-TurboID (SID). TurboID-based biotin ligases biotinylate proximal proteins in the presence of Biotin. The diffused red circle indicated the biotinylation range of TurboID (∼10 nm). (C) Localization of biotinlylated proteins in the presence and absence of exogenously supplemented biotin (50 μM) in mTb-CENP-T cells. (D) Identified kinetochore components and associated factors in the whole-cell extract (WCE), soluble (Sol) and chromatin (Chr) fraction following proximity labeling and IP mass spectrometry (MS) from mTb-NLS, mTb-CENP-T, CENP-T-mTb and SID-CTW. Total spectral counts and exponentially modified protein abundance index (emPAI) values are plot. (E) Western blot analysis of proteins biotinylated by mTb, mTb-NLS, mTb-CT and CT-mTb. Cells were treated with biotin (50 μM) for 6 h, cell fractionated, and biotin affinity pull-down (AP) was performed. (F) Selected top Gene Ontology (GO) analysis terms for the hits described in Figure S4G and S4H, corresponding to the TurboID experiment described in (D). (G) Schematic of strategy used to narrow down TurboID identified hits. Following the AP-MS identification of biotinylated proteins from mTb, mTb-NLS, mTb-CENP-T, CENP-T-mTb and SID-CTW cells, hits were classified as those proteins which are ≥ 3-fold enriched in the CENP-T mTb or CENP-T-W SID over mTb-NLS samples with at least 2 identified spectra. Hits were then considered for a secondary screen if they were identified in ≥ 2 AP-MS samples among the cell fractions of a single cell line or similar fractions across cell lines resulting in 342 proteins hits in total. Following the prediction of essential genes, expected kinetochore and chromosome segregation related factors, and hits with low enrichment were excluded. A single top protein from a well-defined complex was considered. (H) Schematic of Cas13-based knockdown of target mRNA. CasRx from *Ruminococcus flavefaciens* strain XPD3002, as previously described in Konermann, S. *et al.^98^* was used. (I) Schematic of the Cas13-based secondary microscopy screen. For rapid and efficient knockdown of the target mRNA, Cas13-GFP plasmid containing CasRx, 3x pre-guide array and the GFP-reporter was transfected into a strain with stable integration of CasRx expressed from the controllable Tetracycline (Tet-ON) promoter. (J) Localization of CENP-T in Tet-On Cas13 cells in the absence or presence of Cas13-GFP plasmid containing non-target (NT), CENP-T or SENP-6 pre-guide array 30 h post transfection. (K) Secondary Cas13-based microscopic screening of hits. (Top) Plot of identified spectral counts and emPAI of hits chosen for the secondary screen. (Bottom) Mitotic mS-SNAP-CENP-T levels across controls and screening hits were quantified, with CENP-T signal in GFP-positive reporter cells normalized to GFP-negative cells. Mean intensity is reported above each column, mean ± SEM, *n* ≥ 6 except for RFC1 = 4 and no GFP-positive cells were obtained for CSNK2B, TOP2A, GAPDH and PAX-5 across multiple independent experiments, not determined (N.D.).

To identify such factors *in situ*, we employed biotin-based proximity labeling. MiniTurboID (mTb)^50^ was fused to either the N- or C-terminus of endogenous CENP-T (mTb-CT or CT-mTb) to capture spatially distinct interactors (Figures 3B, S3F, S3G and Table S1). In parallel, split-TurboID (SID)^51^ was used to profile interactions specifically engaging with assembled CENP-T-W subcomplex by fusing the inactive TurboID halves to CENP-T and CENP-W (Figures 3B, S3G, S3F and Table S1). MiniTurboID fusions showed robust biotin induction (6 h) with minimal basal activity (Figures 3C and S3H). Of the SID configurations, only N′-SID on CENP-T combined with C′-SID on CENP-W (SID-CTW) yielded sustained labeling (24 h, lower enzymatic activity of SID^52^), likely reflecting structural constraints (Figures S3I-S3K). Fractionation into soluble and chromatin pools, along with analysis of whole-cell extracts, preceded streptavidin pull-down and LC-MS/MS, producing a comprehensive *in situ* spatial map of CENP-T and CENP-T-W proximal interactomes (Figures S3J-S3K).

This proximity labeling analysis recovered nearly all CCAN subunits with the exception of CENP-S, CENP-X, and CENP-W, likely reflecting their small size, lysine content, or extensive biotinylation (Figure 3D). CCAN components were strongly enriched in the chromatin fraction, suggesting that CENP-T assembly into the CCAN occurs predominantly on chromatin rather than in a soluble intermediate, in contrast to previous reports (Figures 3D and 3E).^53^ A notable exception was the CENP-O complex, which appeared in both fractions. The Mis18 complex-an essential regulator of centromere identity and CENP-A deposition^54–58^ - was also highly enriched, suggesting expanded activity at DT40 kinetochores. As expected, outer kinetochore components were more enriched in the mTb-CT fusion relative to CT-mTb (Figure 3D).

Transient interactors, including chromosome passenger complex (CPC) components and spindle assembly checkpoint proteins, were also detected (Figure 3D). The desumoylase, SENP6, a recently identified CCAN regulator was also recovered.^7,8,59,60^ We noted modest enrichment of SSRP1-but not SPT16 -of the FACT complex, which was previously reported to interact with the CCAN^61,62^ including CENP-T^63^ (Table S1). However, conditional depletion of SSRP1 showed no effect on CENP-T complex stability in this system (Figures S3L-S3N).

Collectively, these results define the chromatin and soluble environments of CENP-T, highlighting a largely chromatin-restricted mode of CCAN assembly while revealing a potential distinct soluble connection to the CENP-O complex. This framework provides a mechanistic entry point to identify factors that functionally safeguard CENP-T stability.

### Cas13 screening identifies regulators of CENP-T stability

To identify candidate regulators of CENP-T based on the proximity analysis, we performed gene ontology (GO) analysis on the proximity-labeling datasets. This analysis highlighted strong enrichment for components of the Ada-Two-A-containing (ATAC) complex, which share a catalytic core with the SAGA complex (Figures 3F and S4A).^64^ However, mAID2-mediated^65^ depletion of the ATAC subunits, Yeats2 and ZZZ3, did not alter CENP-T levels (Figures S4B and S4C). Further, histone chaperones identified from our proximity analysis failed to efficiently recruit CENP-T to the LacO array (Figures S1F, S4D-F), suggesting that they are not sufficient to deposit CENP-T.

To refine candidates for a secondary analysis, we selected proteins detected in at least two independent IP-MS experiments and included top hits from each fraction, yielding 342 proteins (Figures 3G, S4G and S4H). Using human essentiality datasets,^66^ reasoning that factors required to maintain CENP-T are likely to be essential, and excluding known chromosome segregation factors, ribosomal proteins, and redundant complex members, we defined a shortlist of 80 candidates (Figure 3G).

To test their functional contributions, we developed a Cas13-based ^98^ microscopy strategy to assay CENP-T stability upon conditional knockdown (cKD) of each candidate, using a pre-guide array (three guides per gene) expressed from a single pCas13-GFP plasmid (Figures 3H and 3I). We quantified CENP-T intensity in GFP-positive cells, with GFP-negative cells serving as internal controls (Figure 3J). As expected, depletion of CENP-T or the CCAN component CENP-I markedly reduced CENP-T signal intensity (reduced by ∼65% and ∼80% relative to controls, respectively; Figures 3J and 3K, S4I).^34^ Depletion of CENP-S or its binding partner CENP-X reduced CENP-T by 30-40%, whereas CENP-U depletion had no effect, consistent with previous reports (Figure 3K).^67^

A notable hit was the desumoylase SENP6, whose depletion caused a profound ∼80% reduction in CENP-T levels (Figure 3K), highlighting a deeply conserved role for SUMO-dependent quality control in CCAN maintenance across vertebrates.^7,8,59,68^ Additional candidates produced moderate (∼20%) decreases (INTS13, ORC2, KIF2C, RBM23), whereas depletion of AATF, POLA1, RFC1, and IWS1 modestly increased CENP-T levels (∼15%). Collectively, these results reveal multiple candidate regulators that modulate CENP-T stability.

### CENP-S-X and the CENP-O complex act synergistically to maintain CENP-T stability

Our initial Cas13 analysis revealed several genes whose individual depletion caused only mild reductions in CENP-T, suggesting that parallel pathways likely safeguard CENP-T stability at kinetochores. To uncover such redundancy, we repeated the Cas13 analysis in a *cenp-s^1/1^* background, which partially destabilizes CENP-T while retaining viability (Figure 2B-2D). In this sensitized setting, Cas13 cKD of selected hits, including EHMT1, CFAP20, and multiple CENP-O complex subunits (CENP-U, -O, -R), produced a synergistic loss of CENP-T (Figure 4A).

**Figure 4.**
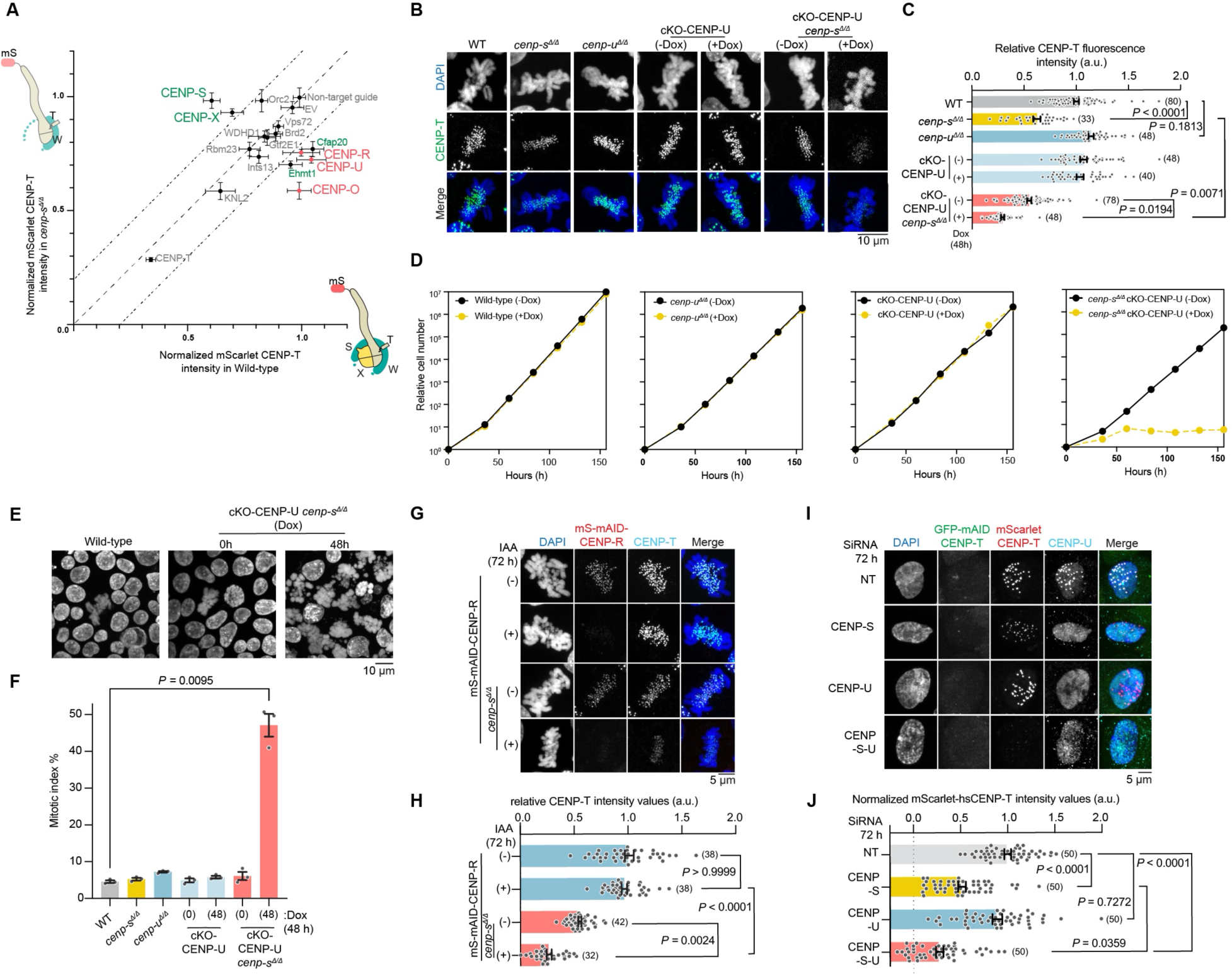
CENP-S-X and the CENP-O complexes coordinately maintain CENP-T stability at kinetochores. (A) Quantified mScarlet-CENP-T levels in mitotic cells following Cas13-based knockdown of selected genes in wild-type vs *cenp-s^1/1^*. Mitotic CENP-T levels across controls and screening hits were quantified, with CENP-T signal in GFP-positive reporter cells normalized to GFP-negative cells. Mean ± SEM, *n* ≥ 7. (B and C) Localization of CENP-T in WT (CL18), *cenp-s^1/1^, cenp-u^1/1^*, CENP-U expressed under the Tet-Off promoter (cKO-CENP-U) and cKO-CENP-U *cenp-s^1/1^* cultured with or without doxycycline (Dox). CENP-T signal in mitotic cells was quantified and internally normalized to co-mixed reference cells in (C). Statistical analysis was performed with Kruskal–Wallis one-way analysis followed by Dunn’s multiple comparison test, number (*n*) of mitotic cells quantified is labelled in parenthesis, mean ± SEM, *P* values are reported as indicated. (D) Growth of WT, *cenp-s^1/1^, cenp-u^1/1^*, cKO-CENP-U and cKO-CENP-U *cenp-s^1/1^* cells with or without Dox addition. The cell numbers were normalized to those at 0 h for each cell line. (E and F) Determination of mitotic index in WT, *cenp-s^1/1^, cenp-u^1/1^*, cKO-CENP-U and cKO-CENP-U *cenp-s^1/1^* cells with or without Dox addition. Representative images of DAPI-stained nuclei are shown in (E) for WT and cKO-CENP-U *cenp-s^1/1^* cells with or without Dox addition. Mitotic index was quantified in (F). Statistical analysis was performed with Kruskal–Wallis one-way analysis followed by Dunn’s multiple comparison test, mean ± SD, *n* = 3, > 1500 cells, *P* values are reported as indicated. (G and H) Localization of CENP-T in mS-mini-auxin-inducible degron (mAID)-CENP-R and mS-mAID-CENP-R *cenp-s^1/1^* with and without Indole-3-acetic acid (IAA). CENP-T signal in mitotic cells was quantified and internally normalized to co-mixed reference cells in (H). Statistical analysis was performed with Kruskal–Wallis one-way analysis followed by Dunn’s multiple comparison test, number (*n*) of mitotic cells quantified is labelled in parenthesis, mean ± SEM, *P* values are reported as indicated. (I and J) Localization of CENP-T in human RPE-1 cells following treatment of SiRNA towards control (non-target guide, SiNT), CENP-S (SiCENP-S), CENP-U (siCENP-U) or both CENP-S and CENP-U (SiCENP-S-U). CENP-T intensity in interphase cells were quantified in (J). Statistical analysis was performed with Kruskal–Wallis one-way analysis followed by Dunn’s multiple comparison test, number (*n*) of interphase cells quantified is labelled in parenthesis, mean ± SEM, *P* values are reported as indicated.

Given its substantial architectural position within the CCAN ^13,15,69^ and identification as a close associate of CENP-T in the soluble and chromatin pool, we focused on the CENP-O complex, whose members include CENP-O, -P, -Q, -U, and -R.^67^ Consistent with prior work, CENP-U KO (*cenp-u^1/1^*) alone had minimal impact on CENP-T or cellular fitness (Figures 4B and 4C). ^67^ Strikingly, in *cenp-s^1/1^* cells, CENP-U depletion (Tet-OFF-based conditional KO, cKO) caused an additional ∼50% loss of CENP-T, with the residual signal dropping to ∼20% after 48 h, most acutely in G1 (Figures 4B, 4C, S5A). Although CENP-U depletion alone was tolerated, combined loss of CENP-S and CENP-U triggered a severe growth arrest and elevated mitotic index, revealing a synthetic-lethal relationship interaction driven by the dramatic reduction in CENP-T (Figures 4D-4F).

To determine whether this synergy reflected a broader dependence on the CENP-O complex, we generated mAID-based cKO CENP-O cells (Figures S5B and S5C) and mAID-based cKO CENP-R (S5D and S5E) cells in the *cenp-s^1/1^* background. Depletion of either subunit similarly reduced CENP-T at kinetochores (Figures 4G, 4H, 5C and 5D) and increased the mitotic index (Figures 5G, 5H, S5F and S5G). These findings underscore a conserved functional link between CENP-S-X-dependent stabilization and CENP-O-complex activity. The milder phenotypes of CENP-R and CENP-O likely reflect slower mAID-mediated depletion.

**Figure 5.**
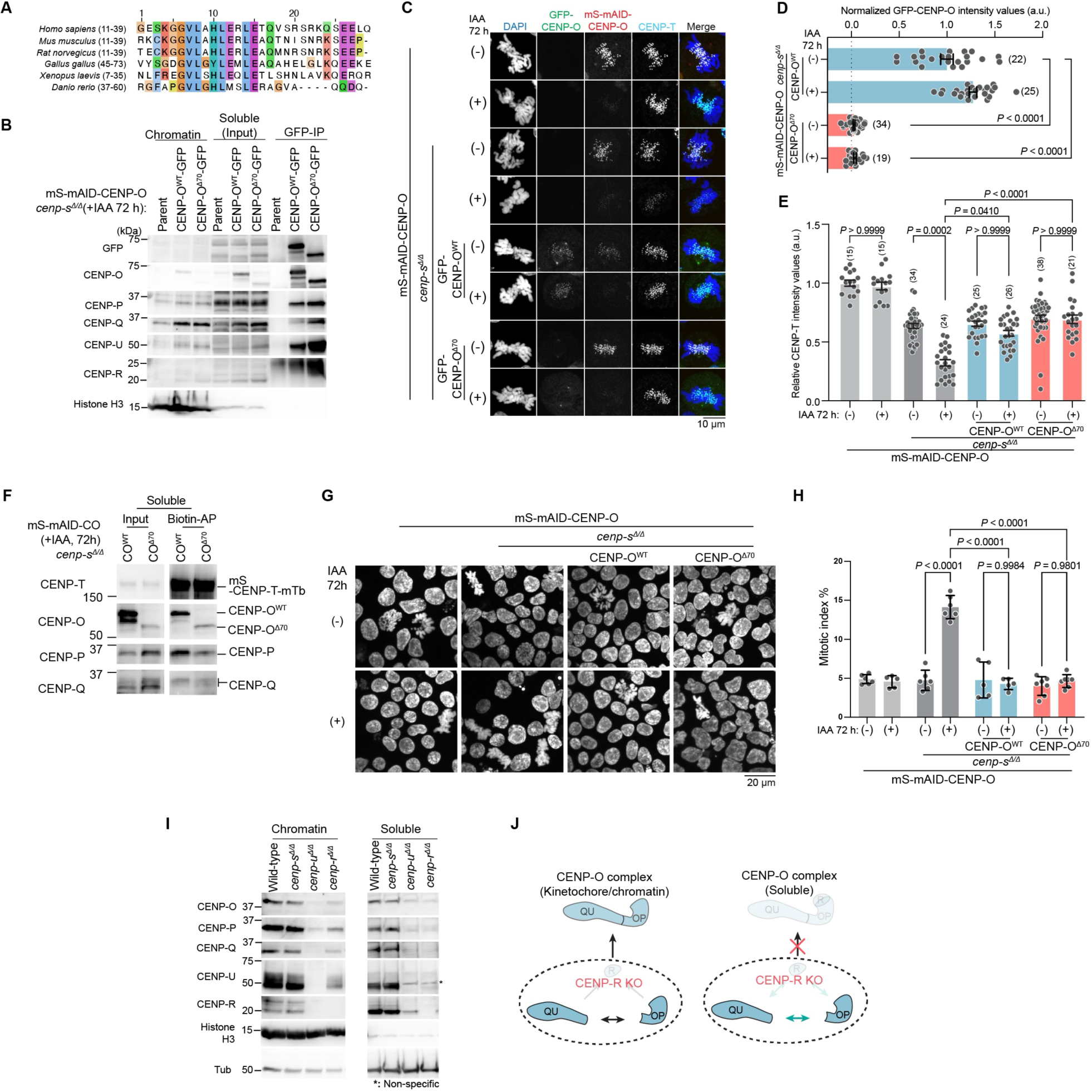
The CENP-O complex regulates CENP-T kinetochore levels via a non-canonical role in the soluble pool. (A) Multiple sequence alignment of the kinetochore targeting region on CENP-O. (B) GFP-IP of CENP-O^WT^ and CENP-O^170^. Following the degradation of mS-mAID-CENP-O, IAA addition 72h, GFP IP of the soluble fraction from the parent (mS-mAID-CENP-O *cenp-s^1/1^*) and the reintegration strains with CENP-O^WT^-GFP or CENP-O^170^-GFP were performed. (C, D and E) Localization of CENP-T in mS-mAID-CENP-O, mS-mAID-CENP-O *cenp-s^1/1^*, and the reintegration strains of CENP-O^WT^-GFP or CENP-O^170^-GFP with and without IAA, 72 h. CENP-O GFP signal in mitotic cells was quantified in (D). CENP-T signal in mitotic cells was quantified and internally normalized to co-mixed reference cells in (E). Statistical analysis was performed with Kruskal–Wallis one-way analysis followed by Dunn’s multiple comparison test in (D and E), number (*n*) of mitotic cells quantified is labelled in parenthesis, mean ± SEM, *P* values are reported as indicated. (F) Western blot analysis of proteins biotinylated by CENP-T-mTb in mS-mAID-CENP-O *cenp-s^1/1^* with CENP-O^WT^-GFP or CENP-O^170^-GFP reintegrated. Cells were grown in the presence of IAA for 72h and treated with biotin (50 μM) for the final 6 h, cell fractionated, and biotin affinity pull-down (AP) was performed from the soluble fraction. *n* = 2. (G and H) Mitotic index of mS-mAID-CENP-O, mS-mAID-CENP-O *cenp-s^1/1^*, and the reintegration strains of CENP-O^WT^-GFP or CENP-O^170^-GFP with and without IAA, 72 h. Representative images of DAPI-stained nuclei are shown. Mitotic index was quantified in (G). Statistical analysis was performed with Kruskal–Wallis one-way analysis followed by Dunn’s multiple comparison test, mean ± SD, *n* ≥ 5, > 1000 cells, *P* values are reported as indicated. (I) Protein levels of CENP-O complex components in the soluble and chromatin pools from wild-type (CL18), *cenp-s^1/1^*, *cenp-u^1/1^* and *cenp-r^1/1^* cells. (J) Schematic depicting how CENP-R differentially contributes to CENP-O complex stability at the kinetochore/chromatin and within the soluble pool.

Finally, to test whether this relationship is conserved in vertebrates, we performed siRNA knockdowns in human RPE-1 cells. As in DT40 cells, single depletion of CENP-U or CENP-O did not alter CENP-T levels at the kinetochore. In contrast, co-depletion of CENP-S and CENP-U significantly reduced kinetochore-bound CENP-T (Figures 4I, 4J, S5H, S5I). These results establish that CENP-S-X and the CENP-O complex constitute parallel, mutually reinforcing pathways that together maintain CENP-T homeostasis and ensure robust kinetochore function.

### A Soluble CENP-O complex regulates CENP-T maintenance at the kinetochore

To define how the CENP-S-X and CENP-O complexes cooperatively maintain CENP-T, we first dissected the CENP-S contribution. Using CENP-S mutants defective in DNA-binding (CENP-S^1C^ and CENP-S^1C-DNA^) or tetramerization (CENP-S^tet^) in cKO CENP-U/*cenp-s^1/1^* cells, we observed strong increases in mitotic index and loss of viability upon CENP-U depletion (Figures S6A and S6B). These phenotypes were shared across all CENP-S mutants, suggesting that DNA-binding by CENP-S is a central determinant of CENP-T stability that acts in parallel with the CENP-O complex.

We next tested the contributions of the CENP-O complex. Although this complex has been linked to Plk1 recruitment through CENP-U ^67,70–72^ and spindle-associated functions through CENP-Q and CENP-R ^69,73,74^, these canonical activities may not account for CENP-T regulation (Figures S7A-S7I). Indeed, blocking Plk1 activity using a Plk1 inhibitor (Bi2536, Figures S7A and S7B) or stably expressing a CENP-U^62A,63A^ mutant that is defective in Plk1-related phosphorylation events ^67,70,71,75,76^ in cKO CENP-U/*cenp-s^1/1^* cells failed to diminish CENP-T. In fact, the CENP-U^62A,63A^ mutant largely restored CENP-T levels, growth, viability and mitotic progression in the sensitized background (Figures S7C-S7G). Likewise, perturbing spindle microtubules with nocodazole did not alter CENP-T localization (Figures S7H and S7I). Together, these findings suggest the presence of an unrecognized activity for the CENP-O complex.

A key clue emerged from our fractionation analyses: the CENP-O complex, but not other CCAN subcomplexes, closely associates with CENP-T-W in the soluble pool (Figures 3D and 3E). To determine whether kinetochore localization is necessary for function, we used a CENP-O^Δ70^ mutant that deletes its N-terminal aa 1-70 region that medicates CENP-O and CENP-I-K interactions (Figure 5A).^13^ This CENP-O^Δ70^ mutant assembles with other CENP-O complex subunits to form a complex (Figure 5B and S7J) but fails to localize to kinetochores (Figures 5C and 5D). Despite this lack of kinetochore localization, CENP-O^Δ70^ underwent robust biotinylation by CENP-T-mTb in the soluble fraction indicating that the CENP-O-CENP-T interaction occurs outside chromatin (Figure 5F). Strikingly, CENP-O^Δ70^ fully rescued CENP-T levels, mitotic index, and viability in the synthetic lethal mAID-based cKO-CENP-O/*cenp-s^1/1^* cells in the presence of IAA (+IAA) (Figures 5C, 5E, 5G, and 5H). Thus, kinetochore localization is dispensable for the essential function of the CENP-O complex in CENP-T maintenance such that the CENP-O complex exerts a soluble, non-canonical role for sustaining CENP-T at kinetochores.

Although CENP-R is dispensable for kinetochore recruitment of other CENP-O components,^67^ its loss in a *cenp-s^1/1^* cells caused synthetic lethality (Figures 4G, 4H, S5F and S5G). Importantly, we find that this phenotype correlated with a selective destabilization of the soluble CENP-O complex, wherein CENP-R KO (*cenp-r^1/1/1^*) cells almost completely lack soluble CENP-O subunits while retaining their chromatin-bound counterparts (Figures 5I and 5J). This contrasts with *cenp-u^1/1^*, where both soluble and chromatin fractions are depleted. These data identify CENP-R as a key stabilizer of the soluble CENP-O complex, ensuring that sufficient complex is available to support CENP-T maintenance.

Taken together, our findings establish a unified model in which CENP-S-mediated DNA-binding and a soluble pool of the CENP-O complex act cooperatively to maintain CENP-T homeostasis. Rather than functioning solely at kinetochores, the CENP-O complex performs a previously unrecognized soluble role that is essential for sustaining CENP-T and ensuring robust kinetochore assembly.

### The CENP-O complex maintains a soluble CENP-T pool to support kinetochore assembly and cell viability

To understand how the CENP-O complex contributes to CENP-T homeostasis, we first assessed the distribution of CENP-T across cellular fractions. As expected, most CENP-T resided on chromatin, with a smaller but distinct soluble pool (Figure 6A). Loss of the CENP-O complex-via CENP-U KO or conditional CENP-U depletion (+Dox), in a WT or sensitized *cenp-s^1/1^* background, led to a selective and pronounced reduction in soluble CENP-T (Figures 6A, 6B, S7K, and S7L). These findings indicate that an association with the CENP-O complex is required to stabilize free, unassembled CENP-T.

**Figure 6.**
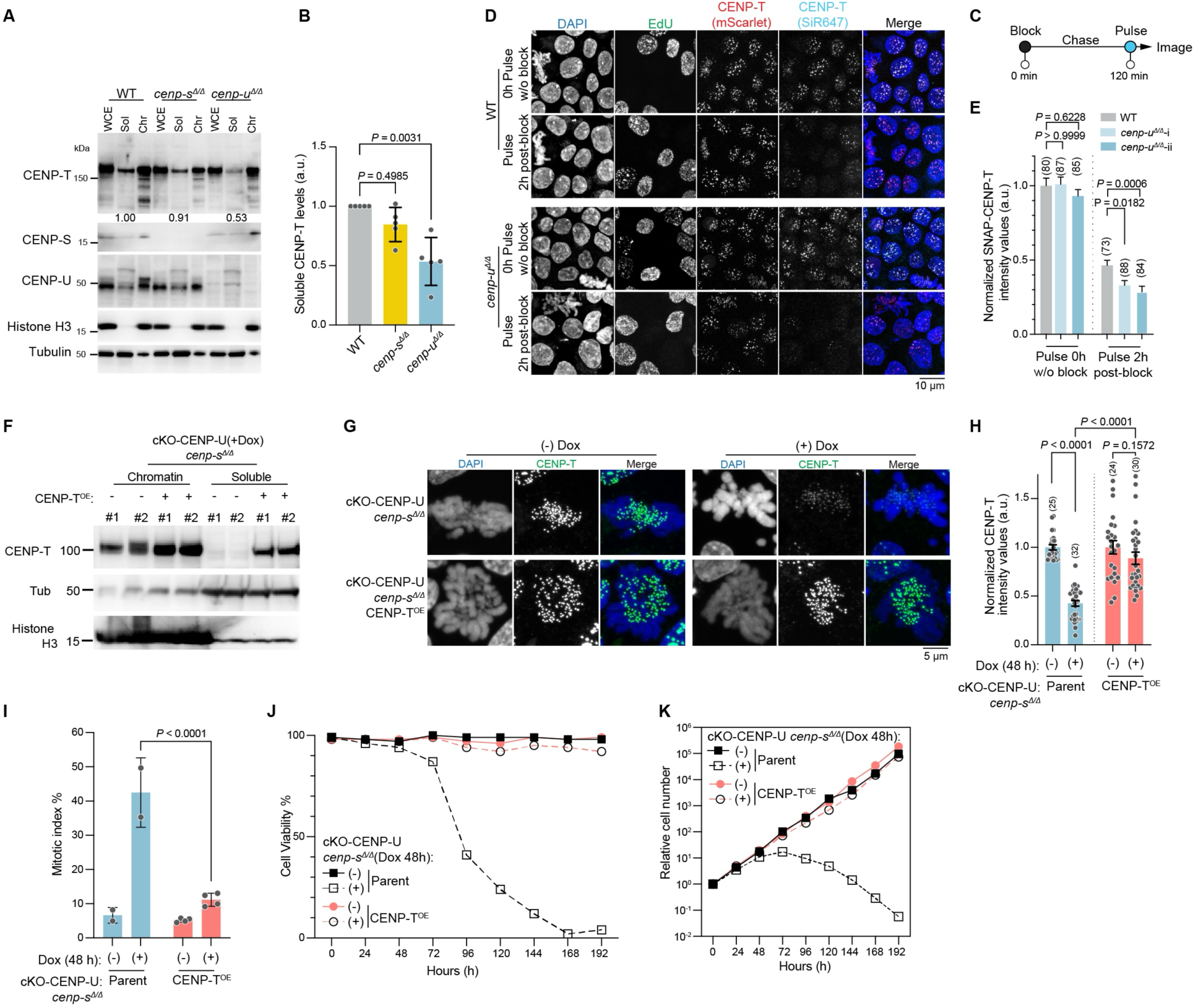
The CENP-O complex preserves the soluble pool of CENP-T, supporting its renewal at the kinetochore. (A and B) CENP-T protein levels across whole-cell extract (WCE), soluble (Sol) and chromatin (Chr) pools in WT (mS-SNAP-CENP-T), *cenp-s^1/1^* and *cenp-u^1/1^* cells. The signal intensity of CENP-T in the soluble pool was quantified in (B). Statistical analysis was performed with Kruskal–Wallis one-way analysis followed by Dunn’s multiple comparison test, mean ± SD, *n* = 5, *P* values are reported as indicated. (C) Schematic of SNAP loading assay. Total cellular mS-SNAP-CENP-T was labelled with a non-fluorescent reagent at 0 h and subsequently washed. Cells were chased for 2 h to allow for new unlabeled protein synthesis and was pulsed with SNAP-Cell 647-SiR, imaged to visualize the localization of newly synthesized protein. (D and E) Localization of SNAP pulsed CENP-T prior to block at 0 h and 2 h post block in wild-type and *cenp-u^1/1^* cells. SNAP labeled CENP-T intensity in G2 were quantified and normalized to its respective 0 h pulse in (E). Statistical analysis was performed with Kruskal–Wallis one-way analysis followed by Dunn’s multiple comparison test, number (*n*) of cells quantified is labelled in parenthesis, mean ± SEM, *P* values are reported as indicated. (F) CENP-T levels in the chromatin-bound and soluble fractions were quantified in parental cKO-CENP-U *cenp-s^1/1^* cells and after integration of the CENP-T^OE^ construct, following 48 h of Dox treatment. (G and H) Localization of CENP-T in the parental cKO-CENP-U *cenp-s^1/1^* cells and following integration of the CENP-T^OE^ construct, determined with or without Dox treatment for 48 h. CENP-T signal in mitotic cells was quantified and internally normalized to co-mixed reference cells (H). Statistical analysis was performed with Kruskal–Wallis one-way analysis followed by Dunn’s multiple comparison test, number (*n*) of mitotic cells quantified is labelled in parenthesis, mean ± SEM, *P* values are reported as indicated. (I, J and K) Mitotic index (F), cell viability (G) and growth (H) of parental cKO-CENP-U *cenp-s^1/1^* cells and after integration of the CENP-T^OE^ construct was quantified with or without Dox treatment for 48 h. Statistical analysis was performed with Kruskal–Wallis one-way analysis followed by Dunn’s multiple comparison test, *n* = 2 and 4 for parent and CENP-T^OE^, respectively in (I). In (K) the cell numbers were normalized to those at 0 h for each cell line.

To determine whether loss of the soluble CENP-O pool also affects CENP-T turnover, we performed SNAP block-pulse-chase assays in *cenp-u^1/1^* cells. We observed a modest but reproducible delay in the replacement of pre-existing CENP-T with newly synthesized protein, despite unchanged total kinetochore levels (Figures 4C, 6C-6E). These data indicate that chromatin-bound CENP-T is stably retained, but its replenishment is slowed when the soluble CENP-O complex is compromised. Thus, regulation of the soluble pool of CENP-T not only sustains steady-state CENP-T abundance but also ensures its timely renewal across cell-cycle turnover.

Together, these results support a two-layered mechanism for CENP-T regulation: 1) the CENP-O complex maintains a sufficient soluble reservoir of CENP-T, to ensure robust replenishment after turnover, 2) CENP-S stabilizes chromatin-bound CENP-T by enhancing its DNA-binding capacity. In the absence of CENP-S, chromatin retention is reduced, and turnover is accelerated. In parallel, without the CENP-O complex-mediated stabilization of soluble CENP-T, this increased turnover cannot be compensated by new assembly, leading to dysregulated CENP-T levels that compromise kinetochore integrity and cell viability.

To directly test this model, we asked whether artificially increasing the soluble CENP-T pool could rescue synthetic lethality. We stably integrated a CMV-driven CENP-T transgene (CENP-T^OE^) into cKO CENP-U/*cenp-s^1/1^* cells. Exogenous expression increased the levels of both soluble CENP-T and chromatin-associated CENP-T in CENP-U-depleted (+Dox) cells, restoring kinetochore localization to near-parental levels (Figures 6F-6H). Importantly, both mitotic index and cell viability were significantly rescued (Figures 6I-6K).

Taken together, these results define a cooperative system in which CENP-S and the CENP-O complex jointly regulate CENP-T homeostasis. The soluble stabilization by the CENP-O complex and the chromatin stabilization by CENP-S-X form an integrated mechanism that is essential for maintaining kinetochore structure and ensuring cell survival (Figure 7).

**Figure 7.**
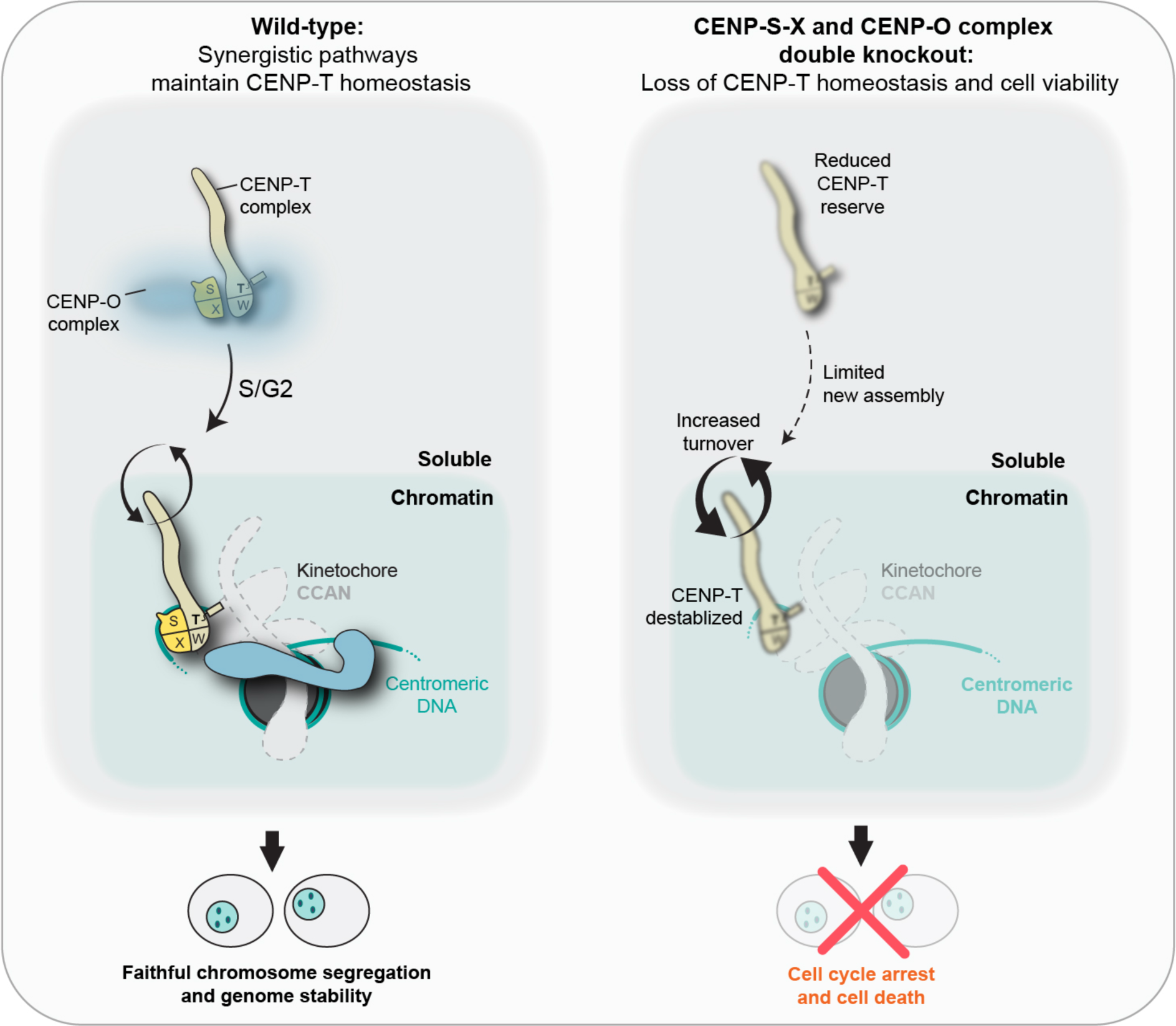
Model for coordinated chromatin stabilization and soluble buffering of CENP-T by CENP-S-X and the CENP-O complex. The CENP-T complex renews each cell cycle, requiring a sufficient soluble pool of CENP-T maintained in part through its close association with the CENP-O complex. Following kinetochore assembly, CENP-T is retained at the kinetochore, until the next turnover cycle, through partial centromeric DNA wrapping driven by the basic C-terminal tail of CENP-S and other basic residues within the complex. Loss of CENP-S-X weakens chromatin retention and increases CENP-T turnover, thereby elevating dependence on the soluble pool, which is further compromised upon loss of the CENP-O complex. Together, destabilization of both soluble and chromatin-bound CENP-T causes collapse of CENP-T homeostasis, resulting in mitotic failure and cell death.

## DISCUSSION

Faithful chromosome segregation requires that kinetochores assemble accurately and be stably maintained throughout each cell cycle. However, the manner in which the kinetochore scaffold combines persistence with the capacity for renewal remained unclear. Here, we reveal that the CENP-T, long regarded as a constitutive scaffold, is in fact dynamically renewed each cycle, and its homeostasis is ensured by two complementary mechanisms mediated by the CENP-S-X and CENP-O complexes (Figure 7). The CENP-S-X complex stabilizes chromatin-incorporated CENP-T through enhanced DNA-binding, whereas the CENP-O complex preserves a soluble reservoir of the CENP-T complex that supports kinetochore assembly (Figure 7). Together, these safeguards explain how kinetochores maintain structural stability while accommodating renewal across generations.

Our turnover analyses demonstrated that both CENP-T and CENP-S undergo rapid replacement during S/G2, followed by stable retention through mitosis (Figures 1A-1F). This behavior contrasts with the inheritance mode of CENP-A,^37,38^ which is deposited in G1 and retained without turnover each cell cycle and indicates that CENP-T complex renewal is a regulated functional requirement rather than incidental. We propose that S/G2 turnover primes the kinetochore for the incorporation of newly synthesized CENP-T-W-S-X complexes each cycle. Consistent with this model, robust CENP-T assembly is observed under conditions of accelerated turnover, such as in *cenp-s^1/1^* cells (Figures 2K-2M). Thus, turnover acts not as a destabilizing force but as a renewal mechanism, ensuring that older complexes are positively replaced with assembly-competent ones needed for a functional kinetochore, supported by soluble buffering and stabilization once incorporated into chromatin. Why does CENP-T, a constitutive chromatin-bound kinetochore subunit, undergo active turnover each cell cycle, in contrast to the more passively inherited CENP-A? Elucidating its physiological significance remains a subject for future research.

The contribution of CENP-S-X to the stabilization process had previously remained unclear. We show that DNA engagement by CENP-S, mediated largely through its conserved basic C-terminal tail, is critical for stabilizing CENP-T at centromeres (Figures 2F and 2G). This interaction likely positions DNA wrapping around the CENP-T complex (Figure 2N), consistent with biochemical evidence that the CENP-T-W-S-X tetramer forms a nucleosome-like particle with strong DNA-binding and supercoiling activity. ^25,35^ Importantly, DNA-binding activity alone is not sufficient for CENP-T stabilization: CENP-S^ΔC^ mutant as part of the CENP-T complex still bound DNA in vitro^25^ but failed to stabilize CENP-T in vivo (Figures 2F and 2G), demonstrating that proper positioning of DNA is also required for CENP-T stabilization. This is in line with reports of CENP-T positioning at the centromere ^78^ and the importance of CENP-S in preventing centromere drift.^79^ In the absence of CENP-S, CENP-T incorporates into chromatin but undergoes accelerated turnover, (Figures 2H-2J), indicating that CENP-S functions as a stabilizer rather than a loader, enhancing DNA engagement to ensure persistence of CENP-T until the next cycle. Supporting these in vivo findings, recent reports suggest that ∼40 bp of DNA wrap the CENP-T-W-S-X tetramer in a nucleosome-like manner, with the CENP-S C-terminal helix becoming ordered upon DNA-binding.^44^ Thus, providing structural confirmation that CENP-S-mediated DNA engagement underlies CENP-T complex’s stable centromeric persistence.

In parallel, the CENP-O complex plays a distinct and complementary role by maintaining soluble CENP-T. Depleting CENP-O components specifically reduces soluble, but not chromatin-bound CENP-T (Figures 6A, 6B, S7H, and S7I). Although this alone does not impair viability, its combination with compromised chromatin-bound CENP-T levels-such as in a CENP-S KO-results in synthetic lethality. This indicates that soluble CENP-T buffering becomes essential when its chromatin retention is compromised. Notably, a kinetochore-localization-defective CENP-O mutant still closely associates with soluble CENP-T and rescues the double mutant (Figures 5C-5H). Demonstrating that the essential function of the CENP-O complex, towards CENP-T complex stability, is not at assembled centromeres but within the soluble assembly pathway.

Within this framework, we also uncovered a previously unappreciated role for CENP-R. Unlike other CENP-O subunits, CENP-R KO mice are viable,^80^ whereas CENP-U KO mice are embryonic lethal. ^81^ In cells, CENP-R kinetochore localization depends on other CENP-O subunits, but not vice versa.^67^ Strikingly, CENP-R depletion collapses soluble CENP-O complex levels (Figure 5I), identifying CENP-R as a stabilizer of soluble complex integrity. Consistent with this role, CENP-R/CENP-S double KO cells are non-viable, highlighting functional specialization within the CENP-O complex.

Comparative analyses suggest evolutionary conservation of this dual-safeguard system. Human cells recapitulate the synthetic reduction of CENP-T observed in chicken DT40 cells upon co-depletion of CENP-S and CENP-U (Figures 4I and 4J). Moreover, the persistence of CENP-R-containing CENP-O complexes across vertebrates^82^ suggests that soluble buffering may be an conserved function, reflecting an evolutionary adaptation of the functionally flexible CENP-O complex.^23^ These safeguards not only preserve stability but may also prevent accumulation of excess free CENP-T, which could otherwise promote ectopic kinetochore formation, consistent with the modest increase in neocentromere formation upon CENP-U loss.^83^ Building on the architectural framework established by high-resolution cryo-EM studies of the kinetochore, our work reveals the regulatory dynamics that preserve this architecture across cell cycles.

Our proximity-labeling analysis was critical in resolving the spatial context of these interactions. Beyond identifying the CENP-O complex as a soluble interactor of CENP-T, this analysis revealed that CCAN assembly occurs directly at centromeres, rather than as pre-assembled modules (Figures 3D and 3E), refining prior models.^53^ Our analysis also identified other potential regulators of CENP-T turnover or incorporation including through histone chaperones (Anp32E and Ing3), transcription regulators (Ints13, IWS1, RBM23, AATF, and Ehmt1), DNA replication-related (Orc2, RFC1 and POLA1) or microtubule-associated factors (KIF2C and CFAP20) (Figure 3K). Future work will clarify how these factors interface with the CENP-O and CENP-S mediated pathways, including the surprisingly close association between CENP-T and the Mis18 complex, a key CENP-A incorporation factor. Additionally, as biotinylation patterns encode proximity geometry, proximity labeling provides a powerful complement to static structural approaches. This may ultimately reveal how kinetochores are oriented and assembled in situ, in addition to mapping dynamic interactions across species.

By defining distinct soluble and chromatin pathways for CENP-T maintenance and renewal, our study provides a conceptual framework for kinetochore homeostasis. The dual-safeguard architecture we describe may represent a broader principle for chromatin-bound machines that must balance durable anchoring with dynamic renewal. Beyond basic biology, our findings raise the possibility that cells under pre-existing kinetochore stress, such as often observed in tumors, may be particularly vulnerable to perturbations that disrupt this balance.

## ACKNOWLEDGMENTS

The authors are very grateful to members of the Fukagawa Laboratory for their inputs and discussions. We also thank R. Fukuoka for their technical assistance, Y. Takenoshita for helping to generate some knockout lines, and Y. Fukui for an initial study on the CENP-O complex. We are grateful to Iain Cheeseman (Massachusetts Institute of Technology) for insightful feedback on the manuscript. The work was supported by JSPS Kakenhi Grants #23K14125 and #25K18405 and to S.S and CREST of JST (JPMJCR21E6), JSPS KAKENHI Grant #22H00408, #23K18113, #24H02281, and #25H00975 to TF.

## AUTHOR CONTRIBUTIONS

S.S and T.F. conceived the study. S.S designed and performed all the experiments. T.F. contributed to the generation of various DT40 cell lines. R.N. performed the mass spectrometry. S.S and T.F. wrote the initial manuscript and all authors provided feedback. S.S and T.F. obtained funding for the study.

## KEY RESOURCES TABLE

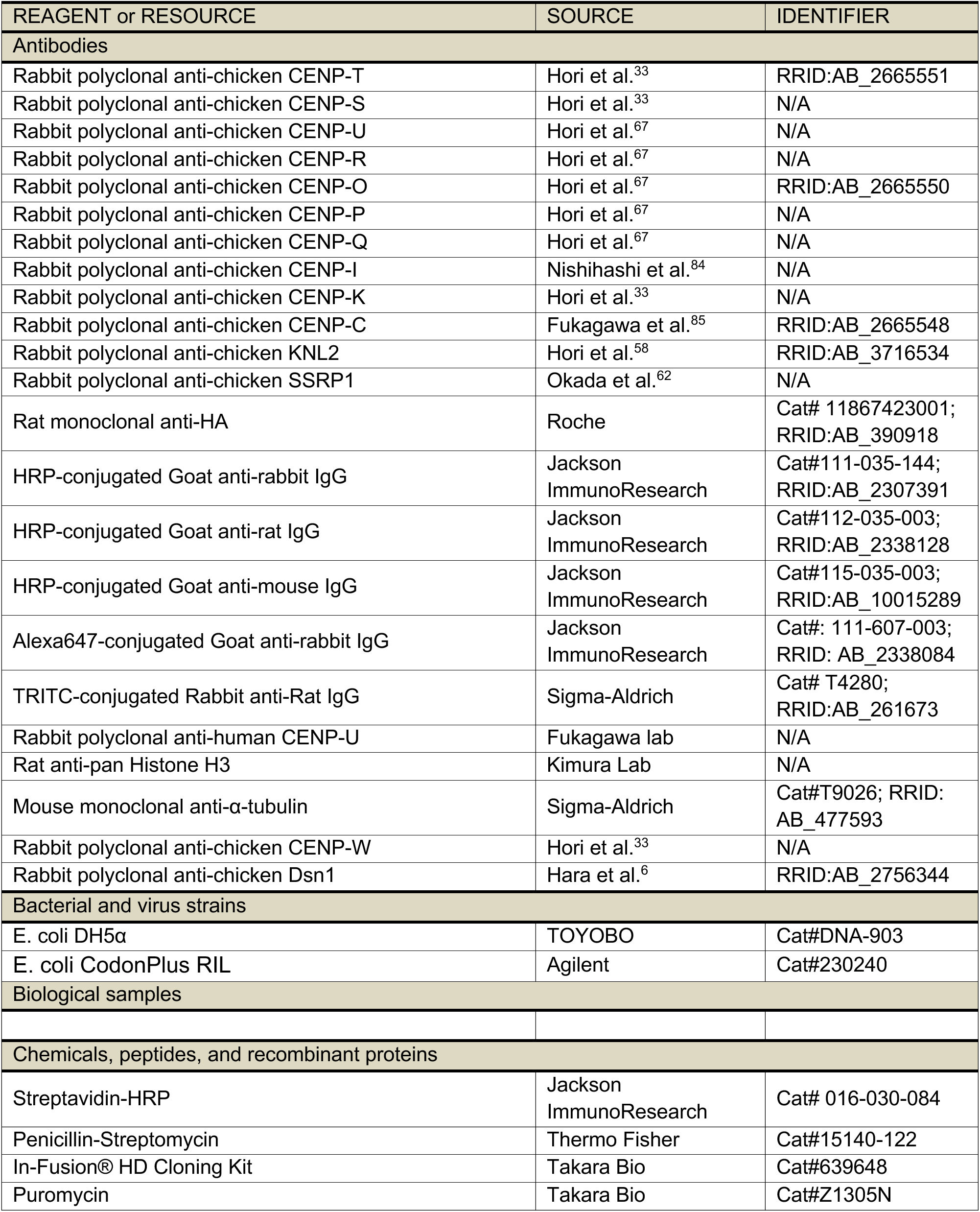
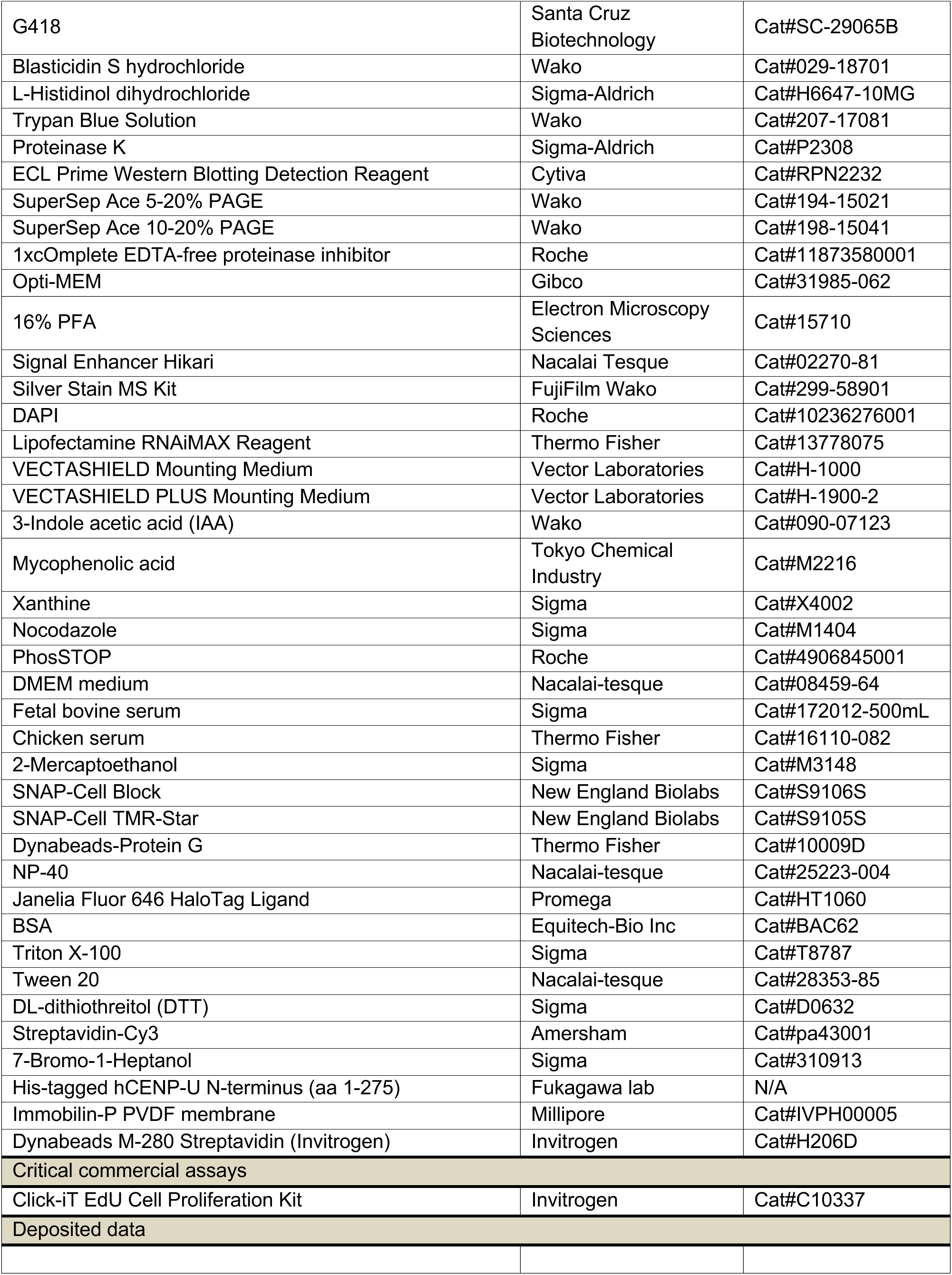
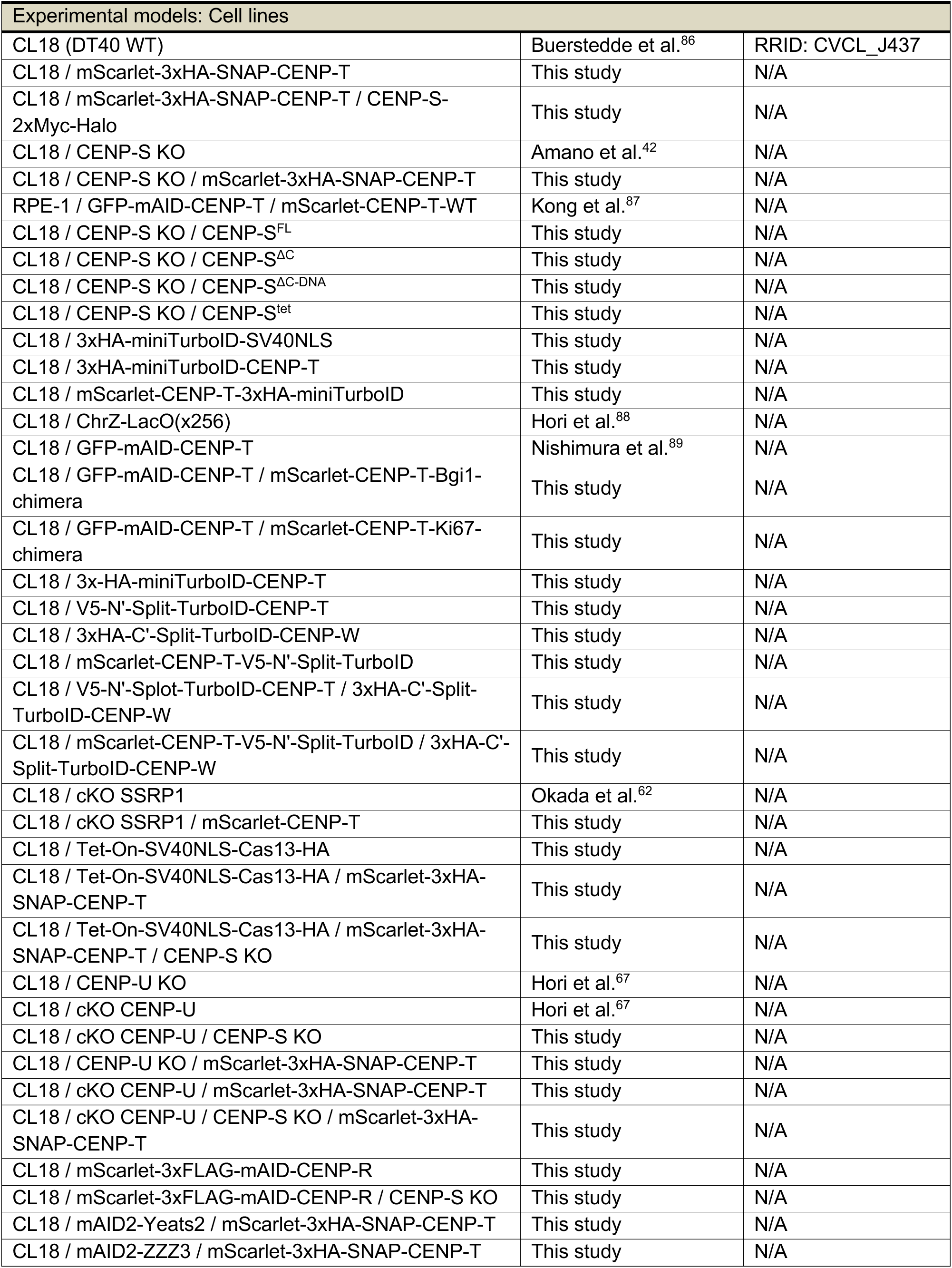
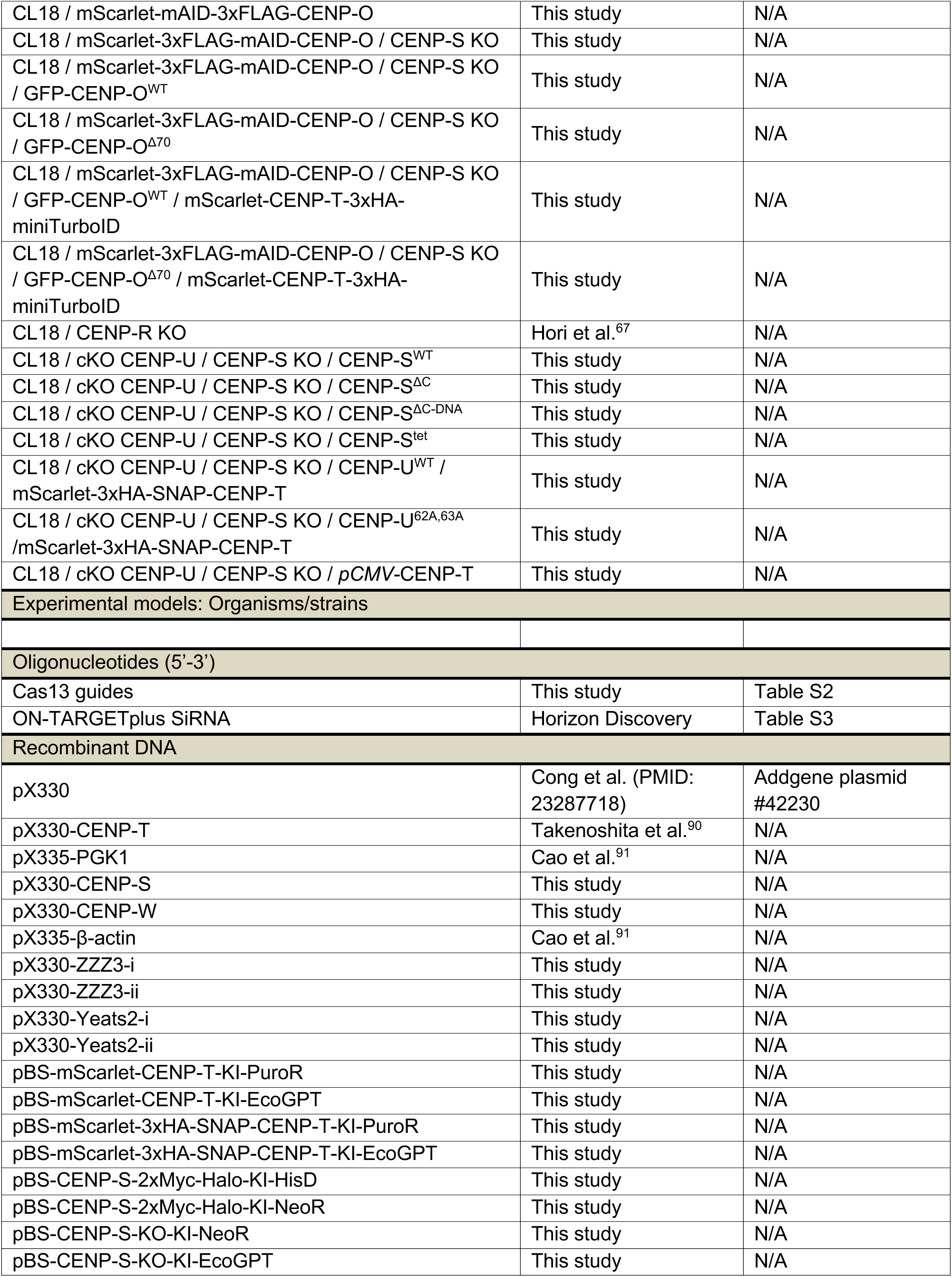
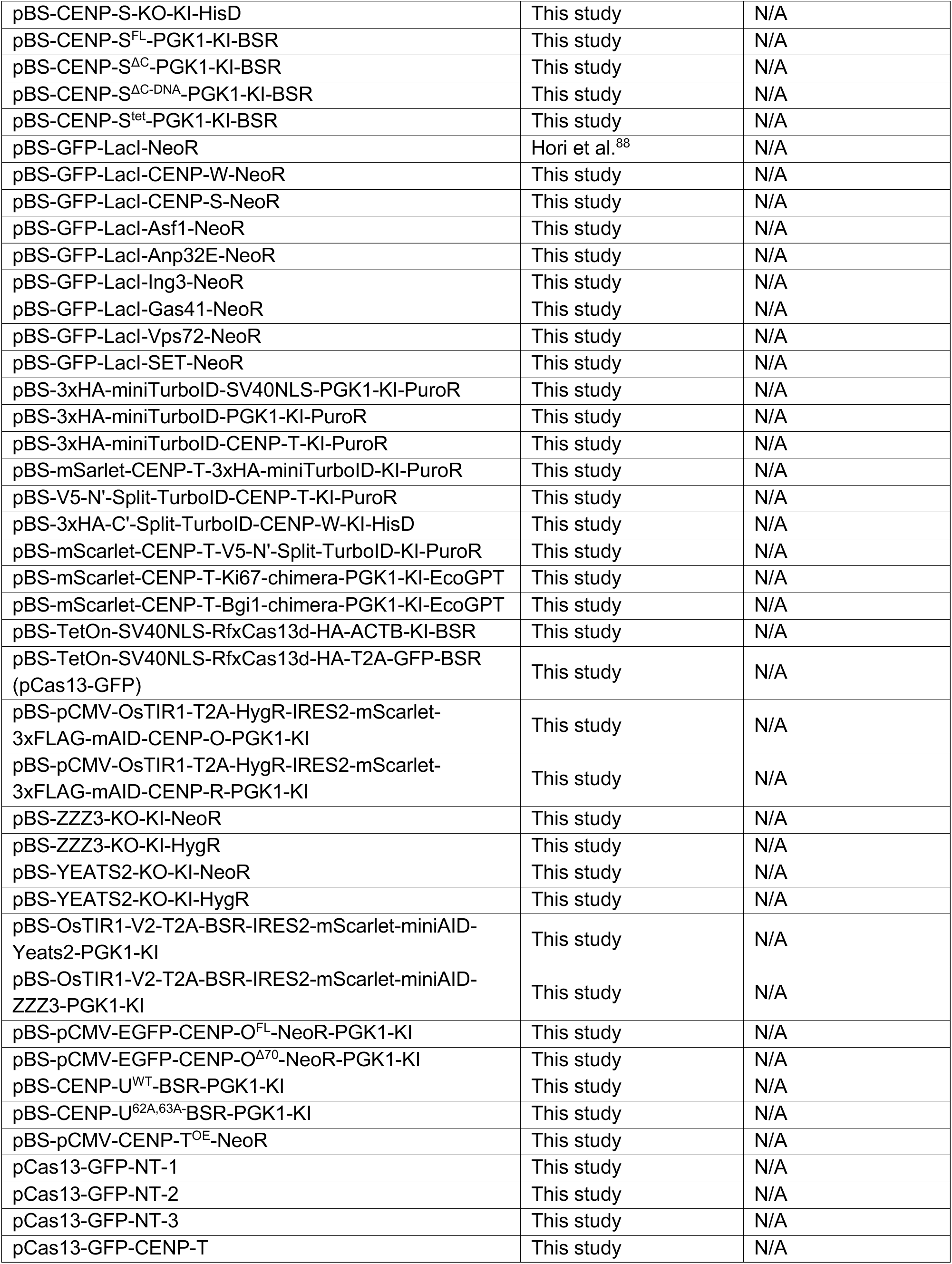
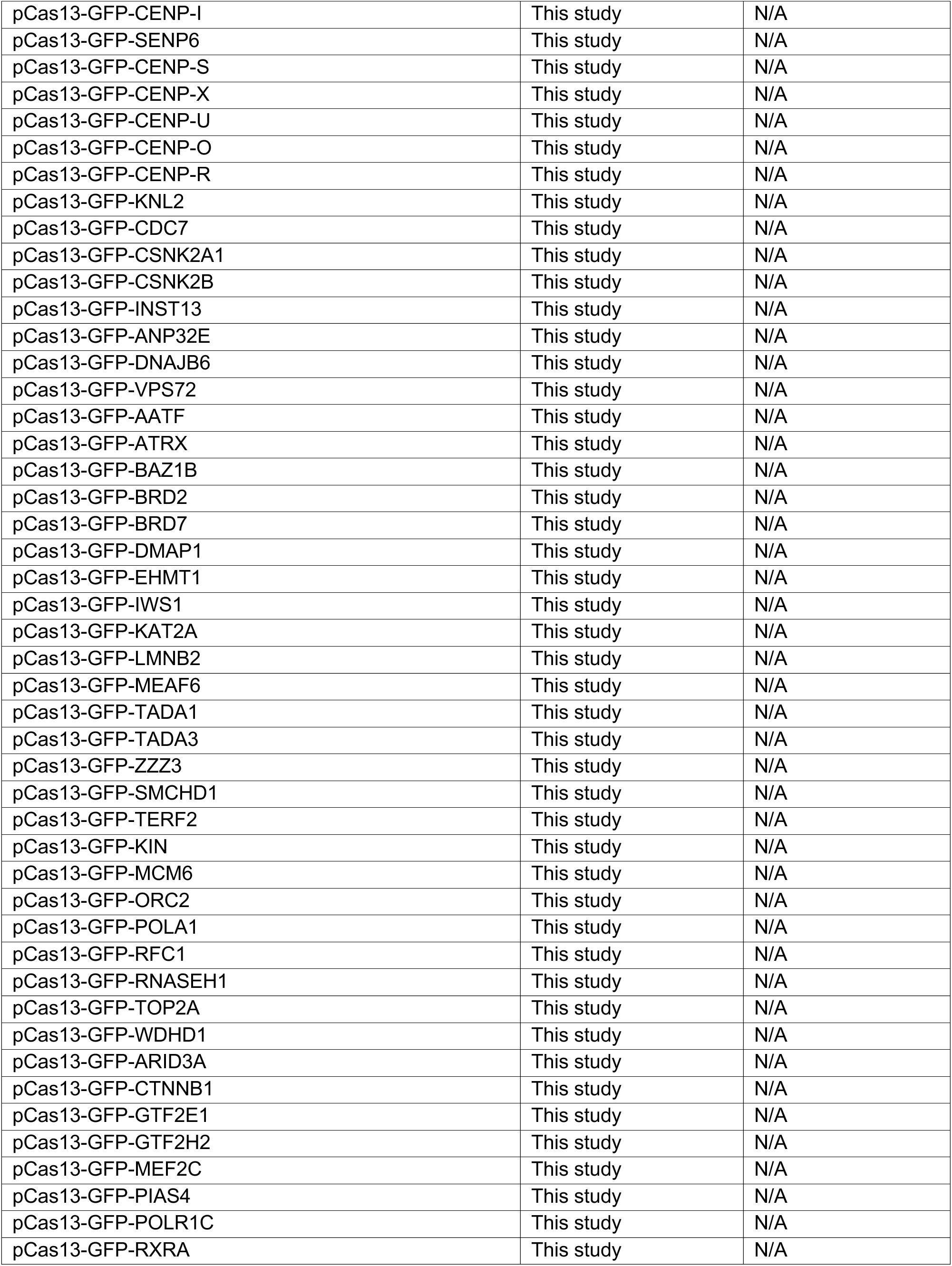
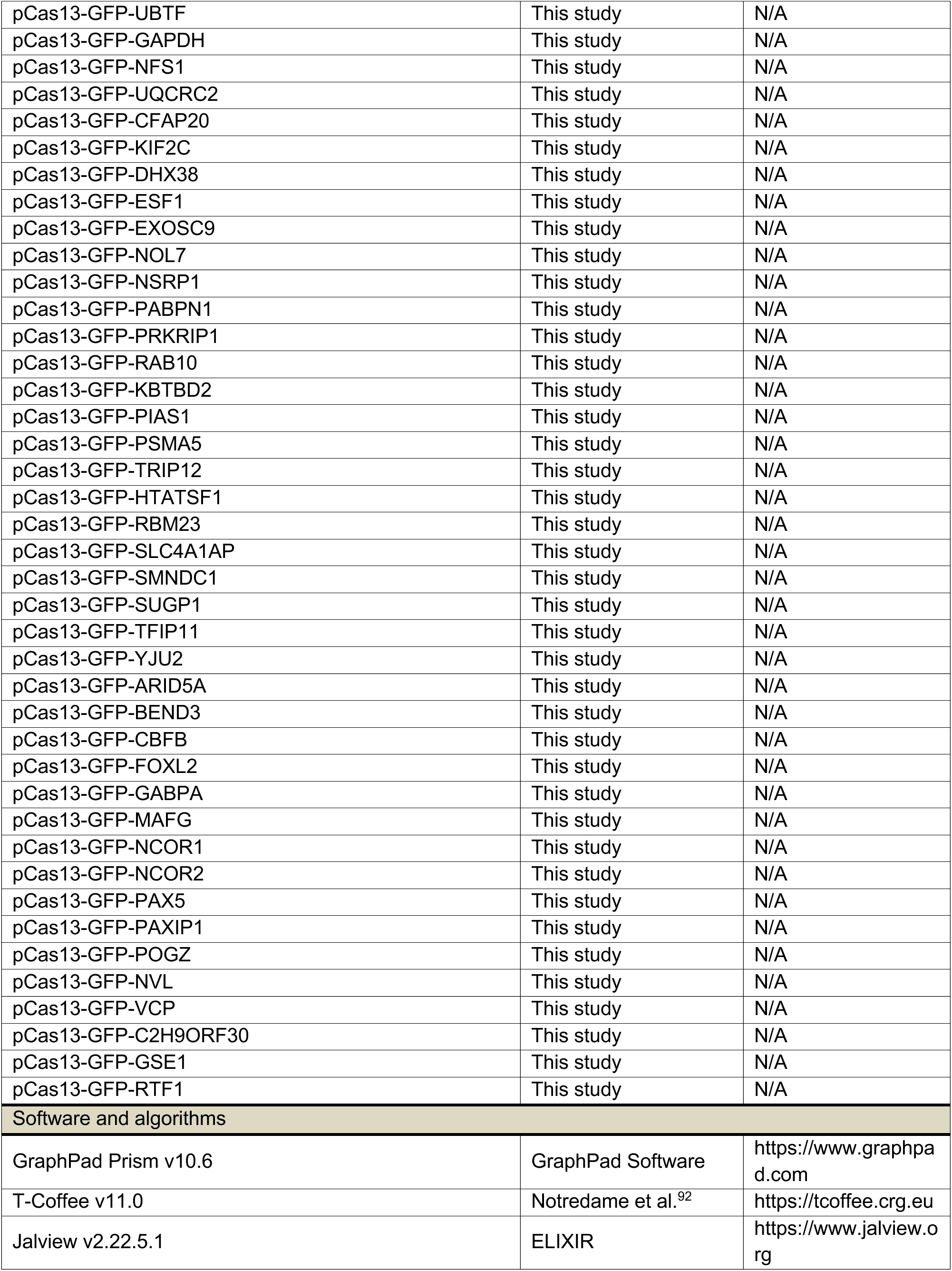
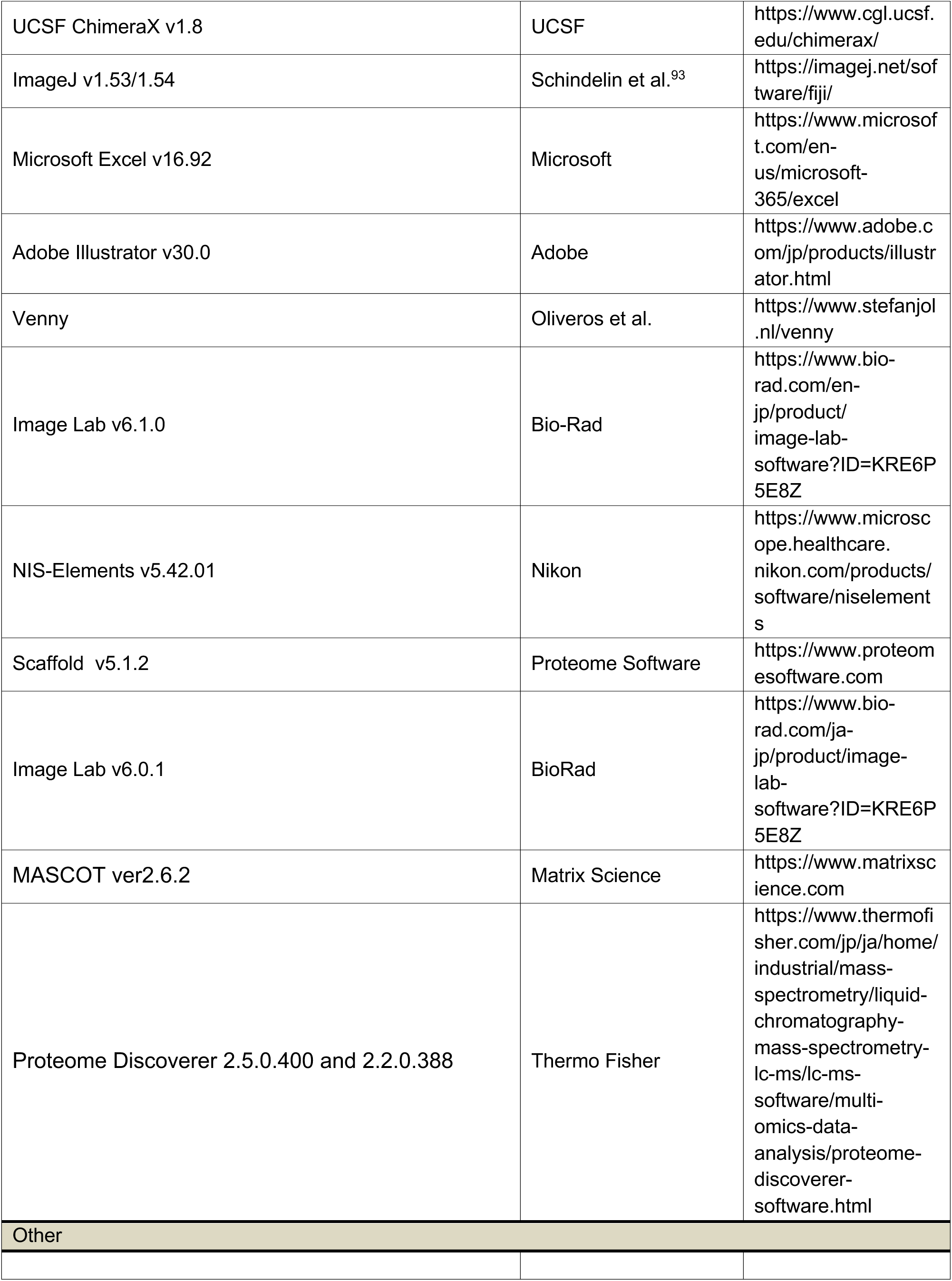

## RESOURCE AVAILABILITY

### Lead contact

Further information and requests for resources and reagents should be directed to and will be fulfilled by the Lead Contact, Tatsuo Fukagawa.

### Materials availability

All unique reagents generated in this study are available from the lead contact with a completed Materials Transfer Agreement.

### Data and code availability

- All data reported in this paper will be shared by the lead contact upon request.
- This paper does not report original code.
- Any additional information required to reanalyze the data reported in this paper is available from the lead contact upon request.

## EXPERIMENTAL MODEL AND STUDY PARTICIPANT DETAILS

### Cell culture

A chicken DT40 cell line CL18 was used as the wild-type (WT) cell.^86^ DT40 cells were cultured in Dulbecco’s modified Eagle’s medium (DMEM, Nacalai-tesque) supplemented with 10% Fetal bovine serum (FBS; Sigma-Aldrich), 1% Chicken serum (Thermo Fisher Scientific), 10 mM 2-Mercaptoethanol (Sigma-Aldrich), Penicillin (100 unit/ml, Thermo Fisher)-Streptomycin (100 mg/ml, Thermo Fisher) at 38.5 °C with 5% CO2. For degradation of mAID-based fusion proteins, cells were treated with 500 μM of indole-3-acetic acid (IAA; Wako). While, for degradation of AID2-based fusion proteins, cells were treated with 1μM 5-Phenyl-IAA (5-Ph-IAA, Sigma-Aldrich). To conditionally knockout CENP-U and SSRP1, cells were cultured in the presence of 2 μg/ml doxycycline (Dox, Sigma-Aldrich).

Human RPE-1 cells were cultured in DMEM (Nacalai-tesque) containing 10% FBS (Sigma-Aldrich) and penicillin (100 unit/mL, Thermo Fisher)-streptomycin (100 μg/mL, Thermo Fisher) at 37 °C with 5% CO2. For degradation of GFP-mAID-CENP-T (RPE-1 cKO-CENP-T), cells were treated with 500 μM of indole-3-acetic acid (IAA; Wako).

All cell lines utilized in the study are listed in the key resource table.

## METHOD DETAILS

### Cell growth and viability analysis

To quantify cell density and monitor growth kinetics, a 50 μL aliquot of DT40 cell culture was harvested and mixed 1:1 with 0.4% (w/v) Trypan Blue solution (Wako). Total cell counts and the proportion of viable versus dead cells were determined using a Countess II Automated Cell Counter (Thermo Fisher Scientific). Relative cell growth was normalized to the initial cell density measured at 0 h for each experimental condition.

### Mitotic Index Determination

For the assessment of mitotic progression, DT40 cells were collected via cytospin and fixed with 3% paraformaldehyde (PFA; Nacalai Tesque) for 10 min at room temperature (RT). Fixed cells were permeabilized with 0.5% NP-40 (Nacalai Tesque) in PBS for 10 min at RT, followed by a brief rinse in PBS. Nuclei were stained with 100 ng/mL DAPI (4′,6-diamidino-2-phenylindole; Sigma-Aldrich) for 15 min at RT, washed, and mounted using VECTASHIELD Mounting Medium (Vector Laboratories).

Nuclear morphology was visualized by fluorescence microscopy to identify mitotic cells characterized by condensed chromosomes (prophase, metaphase, and anaphase). Cells with diffuse, non-condensed chromatin were classified as interphase (non-mitotic). The mitotic index was calculated using the following formula:

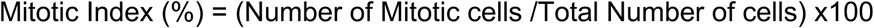

### Plasmid construction and cell line generation

A comprehensive list of cell lines and plasmids used in this study is provided in the key resource table. All plasmids were constructed using the In-Fusion HD Cloning Kit (Takara) and were sequence-verified prior to use.

DT40 cells were transfected using the Neon Transfection System (Thermo Fisher Scientific) with six pulses at 1400 V and 5 ms. Following transfection, cells were plated into 96-well plates to isolate single-cell-derived colonies under appropriate drug selection. Selection was performed using the following concentrations: 0.5 mg/mL puromycin (Takara) for PuroR, 25 mg/mL mycophenolic acid (Wako) plus 125 mg/mL xanthine (Sigma) for EcoGPT, 25 mg/mL blasticidin S hydrochloride (Wako) for BSR, 1 mg/mL L-histidinol for HisD, 1 mg/mL zeocin for BleoR, 2.5 mg/mL hygromycin B for HygR, and 2 mg/mL G418 (Santa Cruz Biotechnology) for NeoR. All engineered cell lines were validated by PCR-based genotyping and further confirmed by immunoblot analysis.

To achieve endogenous replacement and tagged expression under control of native promoters, CRISPR/Cas9-mediated genome editing was employed. For CENP-T tagging donor plasmids encoding mScarlet-CENP-T, mScarlet-3xHA-SNAP-CENP-T, 3xHA-miniTurboID-CENP-T, mScarlet-CENP-T-3xHA-miniTurboID, mScarlet-CENP-T-V5-N′-Split-TurboID, or V5-N′-Split-TurboID-CENP-T were cloned into a pBluescript KS(+) backbone and flanked by ∼1 kb homology arms surrounding the CENP-T start codon. These constructs were co-transfected with pX330-CENP-T.^29^

To replace endogenous CENP-S and express constructs under the endogenous promoters, a donor plasmid encoding CENP-S-2xMyc-Halo was flanked by ∼1 kb homology arms spanning exons 2 and 5 and co-transfected with pX330-CENP-S. sgRNAs targeting CENP-S were designed using Cas-Designer (http://www.rgenome.net/casdesigner/) and cloned into pX330 (Addgene #42330). Complete CENP-S knockout lines were generated by replacing exons 2-5 with NeoR or EcoGPT selection cassettes using homologous recombination.

To replace endogenous CENP-W, a donor plasmid encoding 3xHA-C′-Split-TurboID-CENP-W flanked by ∼1 kb homology arms around the start codon was co-transfected with pX330-CENP-W, containing sgRNAs designed using Cas-Designer.

CENP-T chimera constructs, as well as full-length or mutant versions of CENP-S, CENP-U, and CENP-O, were cloned into a pBluescript backbone containing a ∼2 kb homology region surrounding exon 1 of the PGK1 locus and co-transfected with pX335-PGK1,^91^ which encodes SpCas9 nickase (D10A).^94^ Control miniTurboID strains were generated by integrating plasmids encoding either 3xHA-miniTurboID or 3xHA-miniTurboID-SV40NLS at the PGK1 locus using the same strategy.

Cells expressing RfxCas13d were generated by integrating an SV40NLS-RfxCas13d-HA cassette under a Tet-ON promoter into the β-actin locus using a pBluescript-based donor containing ∼1 kb homology arms and co-transfection with pX335-β-actin.^91^

For mAID2-based conditional knockouts of YEATS2 and ZZZ3, donor plasmids encoding mScarlet-mAID2-Yeats2 or mScarlet-mAID2-ZZZ3, together with OsTIR1-V2, were integrated into the β-actin locus. Subsequent deletion of endogenous YEATS2 alleles was performed using NeoR and HygR selection cassettes with sgRNAs cloned into pX330-Yeats2-i and -ii. ZZZ3 alleles were deleted by CRISPR/Cas9-mediated excision of exon 3 using pX330-ZZZ3-i and -ii.

mAID-based CENP-O and CENP-R conditional knockout lines were generated by integrating OsTIR1 and mScarlet-mAID2-3xFLAG-CENP-O or CENP-R constructs at the PGK1 locus in the respective knockout backgrounds,^67^ as described previously.

For CENP-T overexpression, full-length CENP-T was cloned under the control of the *CMV* promoter in a pBluescript backbone and integrated at random into the DT40 genome.

### Antibody generation

The human CENP-U cDNA was PCR-amplified, and the fragment was digested with BamHI and SalI. The restriction enzyme-digested fragment was cloned into the BamHI/XhoI site of pET28a. Recombinant His-tagged hCENP-U N-terminus (amino acids 1-275) was expressed in *Escherichia coli* CodonPlus RIL cells by induction with 1 mM IPTG for 3 h at 37 °C. The *E. coli* cell pellet was then sent to MBL (Medical & Biological Laboratories). MBL performed affinity purification of the recombinant protein and subsequent production of polyclonal antibodies in rabbits.

### Cell fractionation

To separate soluble and chromatin-associated protein fractions, cells were harvested and washed twice with PBS, followed by a wash in TMS buffer (10 mM Tris-HCl pH 7.5, 5 mM MgCl₂, 0.25 M sucrose). Cells were resuspended in TMS-Triton (TMS supplemented with 0.5% Triton X-100 and 0.5 mM dithiothreitol), containing 1x EDTA-free protease inhibitor cocktail (Roche) and 1x PhosSTOP (Roche), and incubated on ice for 10 min.

Lysates were centrifuged at 1,000 x g for 10 min at 4 °C to pellet nuclei and chromatin. The supernatant (Supernatant 1) was collected and kept on ice. The pellet was resuspended in TMS-Triton and centrifuged at 22,000 x g for 40 min at 4 °C. The resulting supernatant (Supernatant 2) was combined with Supernatant 1, supplemented with one-tenth volume of Buffer 2 (110 mM Tris-HCl pH 8.0, 1.65 M NaCl, 2.2 mM EDTA), incubated on ice for 5 min, and centrifuged at 22,000 x g for 10 min at 4 °C. The final supernatant constituted the soluble fraction (cytoplasmic and nucleoplasmic proteins), as confirmed by exclusive recovery of the NLS-fused miniTurboID control in this fraction.

Pellets from both centrifugation steps were pooled to generate the chromatin fraction, which was resuspended in RIPA buffer (50 mM Tris-HCl, 500 mM NaCl, 0.1% SDS, 0.5% sodium deoxycholate, 1% Triton X-100) supplemented with 1x EDTA-free protease inhibitor cocktail and 1x PhosSTOP. Samples were sonicated to shear DNA and solubilize chromatin-bound proteins.

### Immunoprecipitation

For immunoprecipitation experiments shown in Figures 1I and 5B, cells were harvested and fractionated as described above to obtain the soluble fraction. Soluble extracts were incubated with Protein G Dynabeads (Thermo Fisher Scientific) pre-conjugated with the indicated antibodies for 2 h at 4 °C with gentle rotation.

Bead-bound proteins were washed five times with wash buffer (20 mM Tris-HCl pH 8.0, 150 mM NaCl, 1.5 mM MgCl₂, 0.2 mM EDTA, 0.1% Tween-20) supplemented with 1x EDTA-free protease inhibitor cocktail (Roche) and 1x PhosSTOP (Roche). Proteins were eluted in 1x Laemmli sample buffer (LSB), heated at 96 °C for 10 min, and analyzed by immunoblotting.

### Immunoblot analysis

DT40 cells were harvested, washed with cold PBS, and resuspended in LSB at a final concentration of 1 x 10^4^ cells/µL. Lysates were sonicated and heated at 96 °C for 10 min. Equal amounts of lysate (1 x 10^5^ cells per lane) were resolved by SDS-PAGE (SuperSep Ace, 5-20% or 10-20%; Wako) and transferred onto PVDF membranes (Immobilon-P; Merck).

Membranes were incubated with primary antibodies diluted in Signal Enhancer Hikari Solution A (Nacalai Tesque) overnight at 4 °C, followed by three washes in 0.1% TBST (TBS containing 0.1% Tween-20; 5 min each). Secondary antibodies diluted in Signal Enhancer Hikari Solution B (Nacalai Tesque) were applied for 1 h at room temperature. After three additional washes in 0.1% TBST, signals were detected using ECL Prime (GE Healthcare) and imaged on a ChemiDoc Touch system (Bio-Rad). Image processing and quantification were performed using Image Lab v5.2.1 (Bio-Rad).

For membrane stripping and reprobing, membranes were rinsed in 0.1% TBST for 10 min, then incubated in WB Stripping Solution Strong (Nacalai Tesque), pre-heated to 55 °C, for 20-40 min with gentle agitation. Membranes were washed three times with 0.1% TBST (5 min each), blocked in 2.5% (w/v) BSA for 30 min at room temperature, and subsequently incubated with the next primary antibody.

All antibodies used for immunoblotting in this study are listed in the key resource table.

### TurboID-based biotin proximity labeling and affinity purification

Proximity labeling was performed using TurboID-derived systems (miniTurboID and split-TurboID) to identify proteins proximal to CENP-T or the CENP-T-W complex. Cells expressing the respective TurboID fusion proteins were incubated with 50 µM exogenous biotin for 6 h (miniTurboID) or 24 h (split-TurboID) at 38.5 °C, reflecting the lower catalytic activity of the split enzyme.^52^ During this period, TurboID catalyzes the formation of reactive biotin-AMP, resulting in covalent biotinylation of proteins within an approximately 10 nm radius.

Following labeling, cells were washed once and incubated in fresh, pre-warmed medium for 30 min at 38.5 °C to reduce residual biotin. Cells were then washed once with fresh medium and twice with PBS, after which labeling was quenched by placing cells on ice. Cells were lysed directly in RIPA buffer to generate whole-cell extracts (WCE) or subjected to cell fractionation to obtain soluble and chromatin fractions, as described above.

Biotinylated proteins were purified by incubating lysates or fractionated samples with Dynabeads M-280 Streptavidin (Invitrogen) overnight at 4 °C. Beads were washed sequentially twice with RIPA buffer, once each with 1 M KCl, 0.1 M Na₂CO₃, and 2 M urea in 10 mM Tris-HCl (pH 8.0), and finally twice with RIPA buffer. Bound proteins were eluted by heating the beads at 96 °C for 10 min in LSB supplemented with 2 mM biotin.

### Liquid chromatography and Mass spectrometry

Affinity-purified samples were separated on an SDS-PAGE followed by silver staining (FujiFilm Wako). Silver-stained gels were visualized by ChemiDoc Touch imaging system (Bio-Rad). Isolated samples from the stained gel were Trypsin digested.

Samples were subject to nano LC-MS/MS as described previously.^95^ Briefly, after the gel slices were treated with DTT and iodoacetamide, in-gel digestion with 10 μg/ml modified trypsin (Sequencing grade, Promega) was performed at 37 °C for 16 h. The digested peptides were extracted with 1% trifluoroacetic acid and 50% acetonitrile, dried under a vacuum, and dissolved in 2% acetonitrile and 0.1% formic acid. The peptide mixtures were then fractionated by C18 reverse-phase chromatography (3 μm, ID 0.075 Å∼ 150 mm, CERI, ADVANCE UHPLC, AMR Inc.) at a flow rate of 300 nL/min with a linear gradient of 5-35% solvent B over 20 min. The compositions of solvents A and B were 0.1% TFA in water and 100% acetonitrile, respectively. The data-dependent acquisition was performed by a hybrid linear ion trap mass spectrometer (LTQ Orbitrap Velos Pro, ThermoFisher Scientific) with Advanced Captive Spray SOURCE (AMR Inc.). The mass spectrometer was programmed to carry out 11 successive scans. The first scan was performed as a full-MS scan over the range 350-1800 m/z using Orbitrap analyzer at a resolution of 60,000. The second to eleventh scan events were detected by ion trap analyzer with automatic data-dependent MS/MS scans of the top 10 most abundant ion obtained in the first scan. MS/MS spectra were obtained by setting relative collision energy of 35% CID and exclusion time of 20 s for molecules of the same m/z value range.

Using the MASCOT ver2.6.2 search engine in Proteome Discoverer 2.5.0.400 and 2.2.0.388 (ThermoFisher Scientific), the obtained spectra peaks were assigned using the UniProt proteome database for *Gallus gallus* (ID: UP000000539 (18372 gene count; 43712 entries)). Fragment tolerance 0.80 Da (Monoisotropic), parent tolerance 10 PPM (Monoisotropic), fixed modification of +57 on C (Carbamidomethyl), variable modification of +16 on M (oxidation) and +42 on peptide amino terminus (Acetyl), +226 on K (Biotin) and allowing for a maximum of 2 missed cleavages. Scaffold (v5.1.2, Proteome Software) was used to validate MS/MS-based peptide and protein identifications. Peptide identifications were accepted if they exceeded specific database search engine thresholds. Protein identifications were accepted if they contained at least two identified peptides. Proteins that contained similar peptides and could not be differentiated based on MS/MS analysis alone were grouped to satisfy the principles of parsimony. Proteins sharing significant peptide evidence were grouped into clusters. Identified protein hits and their controls can be found in Supplementary Table 1. To relatively quantitate protein abundance obtained within each of the experiments, emPAI (exponentially modified protein abundance index) ^96^ values were determined using Scaffold (v5.1.2, Proteome Software). The higher the emPAI score, the more abundant the protein is in the mixture.

### LacO/LacI tethering assay

LacO/LacI-based tethering assays were performed using a previously established DT40 cell line in which a 256x LacO repeat array is integrated at the p-arm of chromosome Z.^88^ Plasmids based on the pBluescript (pBS) backbone, expressing GFP-LacI-SV40NLS under the control of the CMV promoter, were introduced into LacO-containing cells by transfection. Following transfection, cells were cultured and analyzed at the indicated time points to assess recruitment of mS-CENP-T to the LacO array.

### Fluorescence microscopy and image acquisition

For visualization of fluorescently tagged proteins (GFP or mScarlet) in DT40 and RPE-1 cells, cells were collected, cytospun onto glass slides, fixed with 3% paraformaldehyde (PFA) for 10 min at room temperature (RT), rinsed with PBS, and permeabilized with 0.5% NP-40 in PBS or 10 min at RT. Where immunostaining was not required, DNA was stained with 100 ng/mL DAPI for 15 min at RT, followed by PBS washes and mounting in VECTASHIELD Mounting Medium (Vector Laboratories).

Immunofluorescence was performed to visualize CENP-T (Figures 2G, 3C, 3J, 4B, 4G, 5C) and Cas13-HA (Figure 3J). Cells were cytospun onto glass slides (Matsunami), fixed with 3% PFA for 10 min at RT, rinsed with PBS, and permeabilized with 0.5% NP-40/PBS for 10 min at RT. Samples were blocked in 0.5% BSA/PBS for 5 min at RT and incubated with primary antibodies diluted in blocking buffer for 1 h at 37 °C. After three washes with 0.5% BSA/PBS, samples were incubated with secondary antibodies for 1 h at 37 °C, washed again, and post-fixed with 3% PFA/PBS for 10 min at RT. DNA was stained with 100 ng/mL DAPI for 15 min at RT prior to mounting in VECTASHIELD or VECTASHIELD PLUS mounting medium.

For visualization of biotinylation (Figure 3C), Streptavidin-Cy3 (1:1000) was included together with the secondary antibody during the secondary incubation step.

To control for staining variability, experiments involving quantification of immuno-stained CENP-T signal (Figures 2G, 4B, 4G, and 5C) included co-cytospinning of test samples with an equal number of reference cells.

For localization of CENP-U in RPE-1 cells (Figures 4I, S5H, and S5I), detached cells were cytospun onto glass slides and fixed with 4% PFA in PHEM buffer (60 mM PIPES, 25 mM HEPES, 10 mM EGTA, 2 mM MgCl₂, pH 6.8) for 10 min at RT. Cells were rinsed with PBS, permeabilized with 0.5% Triton X-100/PBS for 10 min at RT, and blocked for 10 min at RT in blocking buffer (3% BSA, 0.1% Triton X-100, 0.1% sodium azide in TBS). Primary antibodies diluted in blocking buffer were applied for 1 h at 37 °C, followed by three washes in PBST (0.1% Triton X-100/PBS). Secondary antibodies were applied for 1 h at 37 °C, followed by three additional PBST washes. DNA was stained with 100 ng/mL DAPI for 15 min at RT before mounting in VECTASHIELD PLUS.

Fluorescence images were acquired using a Nikon Eclipse Ti inverted microscope equipped with a CSU-W1 spinning-disk confocal unit (Yokogawa), controlled by NIS-Elements (v5.42.01), using PlanApo VC 60x/1.40 NA or Lambda 100x/1.45 NA objectives and an ORCA-Fusion BT sCMOS camera (Hamamatsu). Z-stacks were collected at 0.2-0.3 μm intervals.

Maximum-intensity projections were generated using ImageJ (v1.53)^93^ for representative images.

All antibodies used for immunofluorescence in this study are listed in the key resource table.

### SNAP/Halo assays

For pulse-chase assays, cells expressing mS-SNAP-CENP-T or CENP-S-Halo were incubated with 3 µM SNAP-Cell SiR647 (New England Biolabs) or 200 nM HaloTag JF646 (Promega), respectively, in pre-warmed culture medium for 20 min at 38.5 °C. Following the labeling pulse, cells were rinsed twice with fresh, pre-warmed medium to remove unbound dye. For SNAP labeling, cells were additionally incubated in fresh, pre-warmed medium to further reduce residual dye and background signal. Labeled proteins were then chased for the indicated durations prior to analysis.

For SNAP/Halo loading assays, cells expressing mS-SNAP-CENP-T or CENP-S-Halo were first subjected to a blocking step by incubation with non-fluorescent ligands: 1 mM SNAP-Cell Block (bromothenylpteridine, BTP; New England Biolabs) or 100 nM 7-Bromo-1-Heptanol (Sigma-Aldrich), respectively, in pre-warmed culture medium for 30 min at 38.5 °C. Cells were rinsed twice with fresh, pre-warmed medium and subsequently incubated for an additional 30 min at 38.5 °C in fresh medium to remove residual blocking reagents. Cells were rinsed again and incubated at 38.5 °C for the indicated chase durations, followed by pulse labeling with SNAP or Halo reagents as described above.

For SNAP/Halo experiments, S-phase cells were identified by incorporation of 5-ethynyl-2′-deoxyuridine (EdU; Sigma-Aldrich) using the Click-iT EdU labeling kit (Invitrogen). Cells were pulsed with 10 µM EdU for 15 min at 38.5 °C, cytospun, fixed with 3% paraformaldehyde for 10 min at room temperature, and rinsed with PBS. Samples were permeabilized with 0.5% NP-40 in PBS for 10 min at room temperature, rinsed, and EdU incorporation was detected using copper-catalyzed alkyne-azide cycloaddition by incubation with Click-iT reaction cocktail containing azide-labeled Alexa Fluor 488 for 20 min in the dark at room temperature. Nuclei were counterstained with 100 ng/ml Hoechst 33342 in PBS for 15 min, washed with PBS, and mounted using VECTASHIELD or VECTASHIELD PLUS mounting medium (Vector Laboratories).

### siRNA treatment

For siRNA-mediated knockdown, 10 pmol of the indicated siRNA and 3 µL Lipofectamine RNAiMAX (Thermo Fisher Scientific) were each diluted separately in 125 µL Opti-MEM I Reduced Serum Medium (Gibco). The diluted siRNA and RNAiMAX solutions were then mixed at a 1:1 ratio and incubated for 5 min at room temperature to allow complex formation. The resulting 250 µL siRNA-lipid mixture was added to each well of a 6-well plate (IWAKI), followed by seeding of 5-7.5 x 10^4^ cells per well. Cells were incubated for 3 days at 37 °C prior to analysis. The sequences of siRNAs used in this study are listed in Supplementary Table 3.

### Cas13-based microscopy screen

Guide RNAs for each target gene were designed using the Cas13design tool (https://cas13design.nygenome.org).^95^ Multiple guides targeting non-overlapping regions of each mRNA were selected, with a preference for high-ranking guides targeting the 5′ region of the transcript. All Cas13 guides used in the study are tabulated in Supplementary Table 2.

For each target, a 3x pre-guide array was generated by annealing and extension of 110-bp oligonucleotides, in which each pre-guide was positioned downstream of a 36-bp RfxCas13d direct repeat stem-loop sequence (CAAGTAAACCCCTACCAACTGGTCGGGGTTTGAAAC). The resulting 3x pre-guide arrays were cloned under the U6 promoter into a vector backbone containing Tet-On-Cas13-T2A-GFP using the In-Fusion HD cloning kit (Takara), generating target-specific pCas13 plasmids. The pCas13-GFP construct expresses an active Cas13 effector fused to a C-terminal GFP reporter via a self-cleaving P2A peptide under the control of a doxycycline-inducible Tet-On promoter. Non-targeting control arrays (3x NT guides) and positive control arrays targeting CENP-T or CENP-I were included in each screening set. All plasmids were sequence-verified prior to use.

Target-specific pCas13 plasmids were transfected into DT40 cells expressing SV40NLS-Cas13-HA under the control of a Tet-On promoter. Cells were pre-treated with 0.5 µg/mL doxycycline for 24 h to induce Cas13 expression prior to transfection, which was essential for rapid and efficient depletion of positive-control targets. Continued Cas13 expression from the transfected plasmid was required to robustly maintain knockdown. Transfections were performed using the Neon Transfection System (Thermo Fisher Scientific) with six pulses at 1400 V for 5 ms. Cells were incubated for 30 h post-transfection in the presence of 2 µg/mL doxycycline, harvested, cytospun, and counterstained with DAPI as described above.

For each GFP-positive cell (pCas13 plasmid-positive), kinetochore fluorescence intensities of mScarlet-CENP-T were quantified. To control for slide-to-slide variability and imaging differences, each sample was internally normalized by calculating the ratio of the mean kinetochore intensity in GFP-positive cells to that in GFP-negative cells (pCas13 plasmid-negative).

### Sequence alignment and protein structure visualization

Protein sequences of CENP-O were retrieved from the UniProt database and aligned using T-Coffee (v11.0) to generate high-quality multiple sequence alignments. Alignments were visualized and analyzed using Jalview (v2.22.5.1) to assess domain conservation and identify conserved motifs. For structural analyses, corresponding Protein Data Bank (PDB) structures were obtained and visualized using UCSF ChimeraX (v1.8) to examine spatial organization and residue proximity.

## QUANTIFICATION AND STATISTICAL ANALYSIS

Fluorescence intensities of kinetochore-associated proteins and non-kinetochore background signals were quantified from sum-intensity projections of z-stacks using ImageJ (v1.53/1.54). For each cell, the total kinetochore fluorescence signal was corrected by subtracting the area-normalized background intensity measured from adjacent non-kinetochore regions.

Kinetochore immunofluorescence signals of CENP-T were quantified using ImageJ (v1.53/1.54) for Figures 2H, 4C, 4H, and 5D. To control for variability in immunostaining efficiency and imaging conditions, test samples were co-mixed with an equal number of reference cells at the time of cytospin. The mean kinetochore CENP-T intensity of test cells was first normalized to the mean kinetochore intensity of the reference cells. These normalized values were then divided by the mean normalized value of the control condition to obtain relative CENP-T fluorescence intensities.

Densitometric analysis of immunoblots (Figures 6B and S7I) was performed using ImageJ (v1.54). Integrated pixel densities were measured for soluble CENP-T and the corresponding loading control (α-tubulin) following background subtraction. Relative protein abundance was calculated by normalizing CENP-T signals to α-tubulin for each lane, followed by normalization to the mean value of the control condition.

All quantitative analyses were performed using Microsoft Excel (v16.92) and GraphPad Prism (v10.6; GraphPad Software). Graphs were generated and statistical analyses were conducted using GraphPad Prism. Statistical tests, P values, and error bars (standard deviation or standard error of the mean) are specified in the figure legends. All image processing was performed using ImageJ (v1.53/1.54). When brightness and contrast adjustments were applied, identical settings were used across the entire image. Figures and illustrations were assembled using Adobe Illustrator (v30.0; Adobe). Venn diagrams were generated using Venny (https://www.stefanjol.nl/venny).

## Supplementary Information

The article contains 7 Supplementary Figures, 3 Supplementary Tables.

## REFERENCES

1. Musacchio, A., and Desai, A. (2017). A Molecular View of Kinetochore Assembly and Function. Biology (Basel) 6, 5. 10.3390/biology6010005.

2. Hara, M., Ariyoshi, M., Sano, T., Nozawa, R.-S., Shinkai, S., Onami, S., Jansen, I., Hirota, T., and Fukagawa, T. (2023). Centromere/kinetochore is assembled through CENP-C oligomerization. Mol Cell 83, 2188–2205.e13. 10.1016/j.molcel.2023.05.023.

3. Hara, M., and Fukagawa, T. (2017). Critical Foundation of the Kinetochore: The Constitutive Centromere-Associated Network (CCAN). In Progress in molecular and subcellular biology (Progress in Molecular and Subcellular Biology), pp. 29–57. 10.1007/978-3-319-58592-5_2.

4. Akiyoshi, B., Nelson, C.R., Duggan, N., Ceto, S., Ranish, J.A., and Biggins, S. (2013). The Mub1/Ubr2 Ubiquitin Ligase Complex Regulates the Conserved Dsn1 Kinetochore Protein. PLoS Genet 9. 10.1371/journal.pgen.1003216.

5. Watanabe, R., Hara, M., Okumura, E.I., Hervé, S., Fachinetti, D., Ariyoshi, M., and Fukagawa, T. (2019). CDK1-mediated CENP-C phosphorylation modulates CENP-A binding and mitotic kinetochore localization. Journal of Cell Biology 218, 4042–4062. 10.1083/JCB.201907006.

6. Hara, M., Ariyoshi, M., Okumura, E., Hori, T., and Fukagawa, T. (2018). Multiple phosphorylations control recruitment of the KMN network onto kinetochores. Nat Cell Biol 20, 1378–1388. 10.1038/s41556-018-0230-0.

7. Mukhopadhyay, D., Arnaoutov, A., and Dasso, M. (2010). The SUMO protease SENP6 is essential for inner kinetochore assembly. Journal of Cell Biology 188, 681–692. 10.1083/jcb.200909008.

8. Liebelt, F., Jansen, N.S., Kumar, S., Gracheva, E., Claessens, L.A., Verlaan-de Vries, M., Willemstein, E., and Vertegaal, A.C.O. (2019). The poly-SUMO2/3 protease SENP6 enables assembly of the constitutive centromere-associated network by group deSUMOylation. Nat Commun 10, 3987. 10.1038/s41467-019-11773-x.

9. Oegema, K., Desai, A., Rybina, S., Kirkham, M., and Hyman, A.A. (2001). Functional Analysis of Kinetochore Assembly in Caenorhabditis elegans. Journal of Cell Biology 153, 1209–1226. 10.1083/jcb.153.6.1209.

10. Hanahan, D. (2022). Hallmarks of Cancer: New Dimensions. Cancer Discov 12, 31–46. 10.1158/2159-8290.CD-21-1059.

11. de Wolf, B., and Kops, G.J.P.L. (2017). Kinetochore Malfunction in Human Pathologies. In, pp. 69–91. 10.1007/978-3-319-57127-0_4.

12. Pesenti, M.E., Raisch, T., Conti, D., Walstein, K., Hoffmann, I., Vogt, D., Prumbaum, D., Vetter, I.R., Raunser, S., and Musacchio, A. (2022). Structure of the human inner kinetochore CCAN complex and its significance for human centromere organization. Mol Cell. 10.1016/j.molcel.2022.04.027.

13. Yatskevich, S., Muir, K.W., Bellini, D., Zhang, Z., Yang, J., Tischer, T., Predin, M., Dendooven, T., McLaughlin, S.H., and Barford, D. (2022). Structure of the human inner kinetochore bound to a centromeric CENP-A nucleosome. Science (1979) 376, 844–852. 10.1126/science.abn3810.

14. Yatskevich, S., Yang, J., Bellini, D., Zhang, Z., and Barford, D. (2024). Structure of the human outer kinetochore KMN network complex. Nat Struct Mol Biol 31, 874–883. 10.1038/s41594-024-01249-y.

15. Yan, K., Yang, J., Zhang, Z., McLaughlin, S.H., Chang, L., Fasci, D., Ehrenhofer-Murray, A.E., Heck, A.J.R., and Barford, D. (2019). Structure of the inner kinetochore CCAN complex assembled onto a centromeric nucleosome. Nature 574, 278–282. 10.1038/s41586-019-1609-1.

16. Tromer, E.C., van Hooff, J.J.E., Kops, G.J.P.L., and Snel, B. (2019). Mosaic origin of the eukaryotic kinetochore. Proceedings of the National Academy of Sciences 116, 12873–12882. 10.1073/pnas.1821945116.

17. Sridhar, S., Dumbrepatil, A., Sreekumar, L., Sankaranarayanan, S.R., Guin, K., and Sanyal, K. (2017). Centromere and Kinetochore: Essential Components for Chromosome Segregation. Gene Regulation, Epigenetics and Hormone Signaling, 259–288. 10.1002/9783527697274.ch9.

18. Fukagawa, T., and Earnshaw, W.C. (2014). The centromere: Chromatin foundation for the kinetochore machinery. Dev Cell 30, 496–508. 10.1016/j.devcel.2014.08.016.

19. McKinley, K.L., and Cheeseman, I.M. (2016). The molecular basis for centromere identity and function. Nat Rev Mol Cell Biol 17, 16–29. 10.1038/nrm.2015.5.

20. De Rop, V., Padeganeh, A., and Maddox, P.S. (2012). CENP-A: the key player behind centromere identity, propagation, and kinetochore assembly. Chromosoma 121, 527–538. 10.1007/s00412-012-0386-5.

21. Bui, M., Walkiewicz, M.P., Dimitriadis, E.K., and Dalal, Y. (2013). The CENP-A nucleosome. Nucleus 4, 37–42. 10.4161/nucl.23588.

22. Black, B.E., and Cleveland, D.W. (2011). Epigenetic centromere propagation and the nature of CENP-A nucleosomes. Cell 144, 471–479. 10.1016/j.cell.2011.02.002.

23. Sridhar, S., and Fukagawa, T. (2022). Kinetochore Architecture Employs Diverse Linker Strategies Across Evolution. Front Cell Dev Biol 10. 10.3389/fcell.2022.862637.

24. Hara, M., and Fukagawa, T. (2019). Where is the right path heading from the centromere to spindle microtubules? Cell Cycle 18, 1199–1211. 10.1080/15384101.2019.1617008.

25. Nishino, T., Takeuchi, K., Gascoigne, K.E., Suzuki, A., Hori, T., Oyama, T., Morikawa, K., Cheeseman, I.M., and Fukagawa, T. (2012). CENP-T-W-S-X forms a unique centromeric chromatin structure with a histone-like fold. Cell 148, 487–501. 10.1016/j.cell.2011.11.061.

26. Huisin’T Veld, P.J., Jeganathan, S., Petrovic, A., Singh, P., John, J., Krenn, V., Weissmann, F., Bange, T., and Musacchio, A. (2016). Molecular basis of outer kinetochore assembly on CENP-T. Elife 5, e21007. 10.7554/eLife.21007.

27. Nishino, T., Rago, F., Hori, T., Tomii, K., Cheeseman, I.M., and Fukagawa, T. (2013). CENP-T provides a structural platform for outer kinetochore assembly. EMBO J 32, 424–436. 10.1038/emboj.2012.348.

28. Schleiffer, A., Maier, M., Litos, G., Lampert, F., Hornung, P., Mechtler, K., and Westermann, S. (2012). CENP-T proteins are conserved centromere receptors of the Ndc80 complex. Nat Cell Biol 14, 604–613. 10.1038/ncb2493.

29. Takenoshita, Y., Hara, M., and Fukagawa, T. (2022). Recruitment of two Ndc80 complexes via the CENP-T pathway is sufficient for kinetochore functions. Nat Commun.

30. Lara-Gonzalez, P., Pines, J., and Desai, A. (2021). Spindle assembly checkpoint activation and silencing at kinetochores. Semin Cell Dev Biol 117, 86–98. 10.1016/j.semcdb.2021.06.009.

31. Musacchio, A., and Salmon, E.D. (2007). The spindle-assembly checkpoint in space and time. Nat Rev Mol Cell Biol 8, 379–393. 10.1038/nrm2163.

32. Sissoko, G.B., Tarasovetc, E. V., Marescal, O., Grishchuk, E.L., and Cheeseman, I.M. (2024). Higher-order protein assembly controls kinetochore formation. Nat Cell Biol 26, 45–56. 10.1038/s41556-023-01313-7.

33. Hori, T., Amano, M., Suzuki, A., Backer, C.B., Welburn, J.P., Dong, Y., McEwen, B.F., Shang, W.H., Suzuki, E., Okawa, K., et al. (2008). CCAN Makes Multiple Contacts with Centromeric DNA to Provide Distinct Pathways to the Outer Kinetochore. Cell 135, 1039–1052. 10.1016/j.cell.2008.10.019.

34. Singh, T.R., Saro, D., Ali, A.M., Zheng, X.-F., Du, C., Killen, M.W., Sachpatzidis, A., Wahengbam, K., Pierce, A.J., Xiong, Y., et al. (2010). MHF1-MHF2, a Histone-Fold-Containing Protein Complex, Participates in the Fanconi Anemia Pathway via FANCM. Mol Cell 37, 879–886. 10.1016/j.molcel.2010.01.036.

35. Takeuchi, K., Nishino, T., Mayanagi, K., Horikoshi, N., Osakabe, A., Tachiwana, H., Hori, T., Kurumizaka, H., and Fukagawa, T. (2014). The centromeric nucleosome-like CENP–T–W–S–X complex induces positive supercoils into DNA. Nucleic Acids Res 42, 1644–1655. 10.1093/nar/gkt1124.

36. Yatskevich, S., Muir, K.W., Bellini, D., Zhang, Z., Yang, J., Tischer, T., Predin, M., Dendooven, T., McLaughlin, S.H., and Barford, D. (2022). Structure of the human inner kinetochore bound to a centromeric CENP-A nucleosome. bioArxiv, 2022.01.07.475394. 10.1101/2022.01.07.475394.

37. Jansen, L.E.T., Black, B.E., Foltz, D.R., and Cleveland, D.W. (2007). Propagation of centromeric chromatin requires exit from mitosis. Journal of Cell Biology 176, 795–805. 10.1083/jcb.200701066.

38. Shang, W.-H., Hori, T., Westhorpe, F.G., Godek, K.M., Toyoda, A., Misu, S., Monma, N., Ikeo, K., Carroll, C.W., Takami, Y., et al. (2016). Acetylation of histone H4 lysine 5 and 12 is required for CENP-A deposition into centromeres. Nat Commun 7, 13465. 10.1038/ncomms13465.

39. Keppler, A., Gendreizig, S., Gronemeyer, T., Pick, H., Vogel, H., and Johnsson, K. (2003). A general method for the covalent labeling of fusion proteins with small molecules in vivo. Nat Biotechnol 21, 86–89. 10.1038/nbt765.

40. Los, G. V., Encell, L.P., McDougall, M.G., Hartzell, D.D., Karassina, N., Zimprich, C., Wood, M.G., Learish, R., Ohana, R.F., Urh, M., et al. (2008). HaloTag: A novel protein labeling technology for cell imaging and protein analysis. ACS Chem Biol 3, 373–382. 10.1021/cb800025k.

41. Watanabe, R., Hirano, Y., Hara, M., Hiraoka, Y., and Fukagawa, T. (2022). Mobility of kinetochore proteins measured by FRAP analysis in living cells. Chromosome Research, 1–15. 10.1007/s10577-021-09678-x.

42. Amano, M., Suzuki, A., Hori, T., Backer, C., Okawa, K., Cheeseman, I.M., and Fukagawa, T. (2009). The CENP-S complex is essential for the stable assembly of outer kinetochore structure. Journal of Cell Biology 186, 173–182. 10.1083/jcb.200903100.

43. Nishimura, K., and Kanemaki, M.T. (2014). Rapid Depletion of Budding Yeast Proteins via the Fusion of an Auxin-Inducible Degron (AID). 1–16. 10.1002/0471143030.cb2009s64.

44. Yu, C., Muir, K.W., Yang, J., Zhang, Z., McLaughlin, S.H., and Barford, D. (2025). Models for the architecture of the human inner kinetochore CCAN complex on centromeric α-satellite CENP-A nucleosome arrays. Preprint, 10.1101/2025.11.04.686567 10.1101/2025.11.04.686567.

45. McKinley, K.L., Sekulic, N., Guo, L.Y., Tsinman, T., Black, B.E., and Cheeseman, I.M. (2015). The CENP-L-N Complex Forms a Critical Node in an Integrated Meshwork of Interactions at the Centromere-Kinetochore Interface. Mol Cell 60, 886–898. 10.1016/j.molcel.2015.10.027.

46. Basilico, F., Maffini, S., Weir, J.R., Prumbaum, D., Rojas, A.M., Zimniak, T., De Antoni, A., Jeganathan, S., Voss, B., van Gerwen, S., et al. (2014). The pseudo GTPase CENP-M drives human kinetochore assembly. Elife 3, e02978. 10.7554/eLife.02978.

47. Zhang, Z., Bellini, D., and Barford, D. (2020). Crystal structure of the Cenp-HIKHead-TW sub-module of the inner kinetochore CCAN complex. Nucleic Acids Res 48, 11172–11184. 10.1093/nar/gkaa772.

48. Cuylen, S., Blaukopf, C., Politi, A.Z., Muller-Reichert, T., Neumann, B., Poser, I., Ellenberg, J., Hyman, A.A., and Gerlich, D.W. (2016). Ki-67 acts as a biological surfactant to disperse mitotic chromosomes. Nature 535, 308–312. 10.1038/nature18610.

49. Sridhar, S., Hori, T., Nakagawa, R., Fukagawa, T., and Sanyal, K. (2021). Bridgin connects the outer kinetochore to centromeric chromatin. Nat Commun 12, 146. 10.1038/s41467-020-20161-9.

50. Branon, T.C., Bosch, J.A., Sanchez, A.D., Udeshi, N.D., Svinkina, T., Carr, S.A., Feldman, J.L., Perrimon, N., and Ting, A.Y. (2018). Efficient proximity labeling in living cells and organisms with TurboID. Nat Biotechnol 36, 880–898. 10.1038/nbt.4201.

51. Cho, K.F., Branon, T.C., Rajeev, S., Svinkina, T., Udeshi, N.D., Thoudam, T., Kwak, C., Rhee, H.-W., Lee, I.-K., Carr, S.A., et al. (2020). Split-TurboID enables contact-dependent proximity labeling in cells. Proceedings of the National Academy of Sciences 117, 12143–12154. 10.1073/pnas.1919528117.

52. Cho, K.F., Branon, T.C., Udeshi, N.D., Myers, S.A., Carr, S.A., and Ting, A.Y. (2020). Proximity labeling in mammalian cells with TurboID and split-TurboID. Nat Protoc 15, 3971–3999. 10.1038/s41596-020-0399-0.

53. Hoischen, C., Yavas, S., Wohland, T., and Diekmann, S. (2019). CENP-C/H/I/K/M/T/W/N/L and hMis12 but not CENP-S/X participate in complex formation in the nucleoplasm of living human interphase cells outside centromeres. PLoS One 13, 1–26. 10.1371/journal.pone.0192572.

54. Parashara, P., Medina-Pritchard, B., Abad, M.A., Sotelo-Parrilla, P., Thamkachy, R., Grundei, D., Zou, J., Spanos, C., Kumar, C.N., Basquin, C., et al. (2024). PLK1-mediated phosphorylation cascade activates Mis18 complex to ensure centromere inheritance. Science (1979) 385, 1098–1104. 10.1126/science.ado8270.

55. Thamkachy, R., Medina-Pritchard, B., Park, S.H., Chiodi, C.G., Zou, J., de la Torre-Barranco, M., Shimanaka, K., Abad, M.A., Gallego Páramo, C., Feederle, R., et al. (2024). Structural basis for Mis18 complex assembly and its implications for centromere maintenance. EMBO Rep 25, 3348–3372. 10.1038/s44319-024-00183-w.

56. Hayashi, T., Fujita, Y., Iwasaki, O., Adachi, Y., Takahashi, K., and Yanagida, M. (2004). Mis16 and Mis18 are required for CENP-A loading and histone deacetylation at centromeres. Cell 118, 715–729. 10.1016/j.cell.2004.09.002.

57. Jiang, H., Ariyoshi, M., Hori, T., Watanabe, R., Makino, F., Namba, K., and Fukagawa, T. (2023). The cryo-EM structure of the CENP-A nucleosome in complex with ggKNL2. EMBO J 42. 10.15252/embj.2022111965.

58. Hori, T., Shang, W.-H., Hara, M., Ariyoshi, M., Arimura, Y., Fujita, R., Kurumizaka, H., and Fukagawa, T. (2017). Association of M18BP1/KNL2 with CENP-A Nucleosome Is Essential for Centromere Formation in Non-mammalian Vertebrates. Dev Cell 42, 181–189.e3. 10.1016/j.devcel.2017.06.019.

59. Wagner, K., Kunz, K., Piller, T., Tascher, G., Hölper, S., Stehmeier, P., Keiten-Schmitz, J., Schick, M., Keller, U., and Müller, S. (2019). The SUMO Isopeptidase SENP6 Functions as a Rheostat of Chromatin Residency in Genome Maintenance and Chromosome Dynamics. Cell Rep 29, 480–494.e5. 10.1016/j.celrep.2019.08.106.

60. Mitra, S., Bodor, D.L., David, A.F., Abdul-Zani, I., Mata, J.F., Neumann, B., Reither, S., Tischer, C., and Jansen, L.E.T. (2020). Genetic screening identifies a SUMO protease dynamically maintaining centromeric chromatin. Nat Commun 11, 501. 10.1038/s41467-019-14276-x.

61. Schweighofer, J., Mulay, B., Hoffmann, I., Vogt, D., Pesenti, M.E., and Musacchio, A. (2025). Interactions with multiple inner kinetochore proteins determine mitotic localization of FACT. Journal of Cell Biology 224. 10.1083/jcb.202412042.

62. Okada, M., Okawa, K., Isobe, T., and Fukagawa, T. (2009). CENP-H-containing Complex Facilitates Centromere Deposition of CENP-A in Cooperation with FACT and CHD1. Mol Biol Cell 20, 3986–3995. 10.1091/mbc.E09.

63. Prendergast, L., Müller, S., Liu, Y., Huang, H., Dingli, F., Loew, D., Vassias, I., Patel, D.J., Sullivan, K.F., and Almouzni, G. (2016). The CENP-T/-W complex is a binding partner of the histone chaperone FACT. Genes Dev 30, 1313–1326. 10.1101/gad.275073.115.

64. Spedale, G., Timmers, H.Th.M., and Pijnappel, W.W.M.P. (2012). ATAC-king the complexity of SAGA during evolution. Genes Dev 26, 527–541. 10.1101/gad.184705.111.

65. Yesbolatova, A., Saito, Y., Kitamoto, N., Makino-Itou, H., Ajima, R., Nakano, R., Nakaoka, H., Fukui, K., Gamo, K., Tominari, Y., et al. (2020). The auxin-inducible degron 2 technology provides sharp degradation control in yeast, mammalian cells, and mice. Nat Commun 11, 5701. 10.1038/s41467-020-19532-z.

66. Funk, L., Su, K.-C., Ly, J., Feldman, D., Singh, A., Moodie, B., Blainey, P.C., and Cheeseman, I.M. (2022). The phenotypic landscape of essential human genes. Cell 185, 4634–4653.e22. 10.1016/j.cell.2022.10.017.

67. Hori, T., Okada, M., Maenaka, K., and Fukagawa, T. (2008). CENP-O Class Proteins Form a Stable Complex and Are Required for Proper Kinetochore Function. Mol Biol Cell 19, 843–854. 10.1091/mbc.e07-06-0556.

68. Mitra, S., Bodor, D.L., David, A.F., Abdul-Zani, I., Mata, J.F., Neumann, B., Reither, S., Tischer, C., and Jansen, L.E.T. (2020). Genetic screening identifies a SUMO protease dynamically maintaining centromeric chromatin. Nat Commun 11. 10.1038/s41467-019-14276-x.

69. Pesenti, M.E., Prumbaum, D., Auckland, P., Smith, C.M., Faesen, A.C., Petrovic, A., Erent, M., Maffini, S., Pentakota, S., Weir, J.R., et al. (2018). Reconstitution of a 26-Subunit Human Kinetochore Reveals Cooperative Microtubule Binding by CENP-OPQUR and NDC80. Mol Cell 71, 923–939.e10. 10.1016/j.molcel.2018.07.038.

70. Nguyen, A.L., Fadel, M.D., and Cheeseman, I.M. (2021). Differential requirements for the CENP-O complex reveal parallel PLK1 kinetochore recruitment pathways. Mol Biol Cell 32, 712–721. 10.1091/mbc.E20-11-0751.

71. Singh, P., Pesenti, M.E., Maffini, S., Carmignani, S., Hedtfeld, M., Petrovic, A., Srinivasamani, A., Bange, T., and Musacchio, A. (2021). BUB1 and CENP-U, Primed by CDK1, Are the Main PLK1 Kinetochore Receptors in Mitosis. Mol Cell 81, 67–87.e9. 10.1016/j.molcel.2020.10.040.

72. Chen, Q., Zhang, M., Pan, X., Yuan, X., Zhou, L., Yan, L., Zeng, L.-H., Xu, J., Yang, B., Zhang, L., et al. (2021). Bub1 and CENP-U redundantly recruit Plk1 to stabilize kinetochore-microtubule attachments and ensure accurate chromosome segregation. Cell Rep 36, 109740. 10.1016/j.celrep.2021.109740.

73. Sedzro, D.M., Yuan, X., Mullen, M., Ejaz, U., Yang, T., Liu, X., Song, X., Tang, Y.-C., Pan, W., Zou, P., et al. (2022). Phosphorylation of CENP-R by Aurora B regulates kinetochore–microtubule attachment for accurate chromosome segregation. J Mol Cell Biol 14. 10.1093/jmcb/mjac051.

74. Hu, C., Hu, Q., Yang, T., Xu, P., Xiong, F., Wang, X., Wang, C., Jiang, K., Hill, D.L., Xue, L., et al. (2025). Condensation-dependent multivalent interactions of EB1 and CENP-R regulate chromosome oscillations in mitosis. Cell Rep 44, 115560. 10.1016/j.celrep.2025.115560.

75. Kang, Y.H., Park, J.-E., Yu, L.-R., Soung, N.-K., Yun, S.-M., Bang, J.K., Seong, Y.-S., Yu, H., Garfield, S., Veenstra, T.D., et al. (2006). Self-Regulated Plk1 Recruitment to Kinetochores by the Plk1-PBIP1 Interaction Is Critical for Proper Chromosome Segregation. Mol Cell 24, 409–422. 10.1016/j.molcel.2006.10.016.

76. Kang, Y.H., Park, C.H., Kim, T.-S., Soung, N.-K., Bang, J.K., Kim, B.Y., Park, J.-E., and Lee, K.S. (2011). Mammalian Polo-like Kinase 1-dependent Regulation of the PBIP1-CENP-Q Complex at Kinetochores. Journal of Biological Chemistry 286, 19744–19757. 10.1074/jbc.M111.224105.

77. van den Berg, S.J.W., East, S., Mitra, S., and Jansen, L.E.T. (2023). p97/VCP drives turnover of SUMOylated centromeric CCAN proteins and CENP-A. Mol Biol Cell 34. 10.1091/mbc.E23-01-0035.

78. Thakur, J., and Henikoff, S. (2016). CENPT bridges adjacent CENPA nucleosomes on young human α-satellite dimmers. Genome Res 26, 1178–1187. 10.1101/gr.204784.116.

79. Hori, T., Kagawa, N., Toyoda, A., Fujiyama, A., Misu, S., Monma, N., Makino, F., Ikeo, K., and Fukagawa, T. (2017). Constitutive centromere-associated network controls centromere drift in vertebrate cells. Journal of Cell Biology 216, 101–113. 10.1083/jcb.201605001.

80. Okumura, K., Kagawa, N., Saito, M., Yoshizawa, Y., Munakata, H., Isogai, E., Fukagawa, T., and Wakabayashi, Y. (2017). CENP-R acts bilaterally as a tumor suppressor and as an oncogene in the two-stage skin carcinogenesis model. Cancer Sci 108, 2142–2148. 10.1111/cas.13348.

81. Kagawa, N., Hori, T., Hoki, Y., Hosoya, O., Tsutsui, K., Saga, Y., Sado, T., and Fukagawa, T. (2014). The CENP-O complex requirement varies among different cell types. Chromosome Research 22, 293–303. 10.1007/s10577-014-9404-1.

82. Hooff, J.J., Tromer, E., Wijk, L.M., Snel, B., and Kops, G.J. (2017). Evolutionary dynamics of the kinetochore network in eukaryotes as revealed by comparative genomics. EMBO Rep 18, 1559–1571. 10.15252/embr.201744102.

83. Shang, W.-H., Hori, T., Martins, N.M.C., Toyoda, A., Misu, S., Monma, N., Hiratani, I., Maeshima, K., Ikeo, K., Fujiyama, A., et al. (2013). Chromosome Engineering Allows the Efficient Isolation of Vertebrate Neocentromeres. Dev Cell 24, 635–648. 10.1016/j.devcel.2013.02.009.

84. Nishihashi, A., Haraguchi, T., Hiraoka, Y., Ikemura, T., Regnier, V., Dodson, H., Earnshaw, W.C., and Fukagawa, T. (2002). CENP-I Is Essential for Centromere Function in Vertebrate Cells. Dev Cell 2, 463–476. 10.1016/S1534-5807(02)00144-2.

85. Fukagawa, T., Pendon, C., Morris, J., and Brown, W. (1999). CENP-C is necessary but not sufficient to induce formation of a functional centromere. EMBO J 18, 4196–4209. 10.1093/emboj/18.15.4196.

86. Buerstedde, J.M., Reynaud, C.A., Humphries, E.H., Olson, W., Ewert, D.L., and Weill, J.C. (1990). Light chain gene conversion continues at high rate in an ALV-induced cell line. EMBO J 9, 921–927. 10.1002/j.1460-2075.1990.tb08190.x.

87. Kong, W., Hara, M., Tokunaga, Y., Okumura, K., Hirano, Y., Miao, J., Takenoshita, Y., Hashimoto, M., Sasaki, H., Fujimori, T., et al. (2025). CENP-C-Mis12 complex establishes a regulatory loop through Aurora B for chromosome segregation. Life Sci Alliance 8. 10.26508/lsa.202402927.

88. Hori, T., Shang, W.H., Takeuchi, K., and Fukagawa, T. (2013). The CCAN recruits CENP-A to the centromere and forms the structural core for kinetochore assembly. Journal of Cell Biology 200, 45–60. 10.1083/jcb.201210106.

89. Nishimura, K., and Fukagawa, T. (2017). An efficient method to generate conditional knockout cell lines for essential genes by combination of auxin-inducible degron tag and CRISPR/Cas9. Chromosome Research 25, 253–260. 10.1007/s10577-017-9559-7.

90. Takenoshita, Y., Hara, M., Nakagawa, R., Ariyoshi, M., and Fukagawa, T. (2024). Molecular details and phosphoregulation of the CENP-T-Mis12 complex interaction during mitosis in DT40 cells. iScience 27. 10.1016/j.isci.2024.111295.

91. Cao, J.H., Hori, T., Ariyoshi, M., and Fukagawa, T. (2024). Artificial tethering of constitutive centromere-associated network proteins induces CENP-A deposition without Knl2 in DT40 cells. J Cell Sci 137. 10.1242/jcs.261639.

92. Notredame, C., Higgins, D.G., and Heringa, J. (2000). T-coffee: a novel method for fast and accurate multiple sequence alignment 1 1Edited by J. Thornton. J Mol Biol 302, 205–217. 10.1006/jmbi.2000.4042.

93. Schindelin, J., Arganda-Carreras, I., Frise, E., Kaynig, V., Longair, M., Pietzsch, T., Preibisch, S., Rueden, C., Saalfeld, S., Schmid, B., et al. (2012). Fiji: An open-source platform for biological-image analysis. Nat Methods 9, 676–682. 10.1038/nmeth.2019.

94. Cong, L., Ran, F.A., Cox, D., Lin, S., Barretto, R., Habib, N., Hsu, P.D., Wu, X., Jiang, W., Marraffini, L.A., et al. (2013). Multiplex Genome Engineering Using CRISPR/Cas Systems. Science (1979) 339, 819–823. 10.1126/science.1231143.

95. Oya, E., Nakagawa, R., Yoshimura, Y., Tanaka, M., Nishibuchi, G., Machida, S., Shirai, A., Ekwall, K., Kurumizaka, H., Tagami, H., et al. (2019). H3K14 ubiquitylation promotes H3K9 methylation for heterochromatin assembly. EMBO Rep 20. 10.15252/embr.201948111.

96. Wessels, H.H., Méndez-Mancilla, A., Guo, X., Legut, M., Daniloski, Z., and Sanjana, N.E. (2020). Massively parallel Cas13 screens reveal principles for guide RNA design. Nat Biotechnol 38, 722–727. 10.1038/s41587-020-0456-9.

97. Shinoda, K., Tomita, M., and Ishihama, Y. (2009). emPAI Calc-for the estimation of protein abundance from large-scale identification data by liquid chromatography-tandem mass spectrometry. Bioinformatics 26, 576–577. 10.1093/bioinformatics/btp700.

98. Konermann, S., Lotfy, P., Brideau, N.J., Oki, J., Shokhirev, M.N., and Hsu, P.D. (2018). Transcriptome Engineering with RNA-Targeting Type VI-D CRISPR Effectors. Cell 173, 665–676.e14. 10.1016/j.cell.2018.02.033.

