## Supplemental Figures for "Kinetochore Homeostasis is Maintained by Coordinated Chromatin Stabilization and Soluble Buffering"

**or**

**Tatsuo Fukagawa**

[3]Lead contact: **Tatsuo Fukagawa**

**The PDF file includes:**

Figures S1 to S7

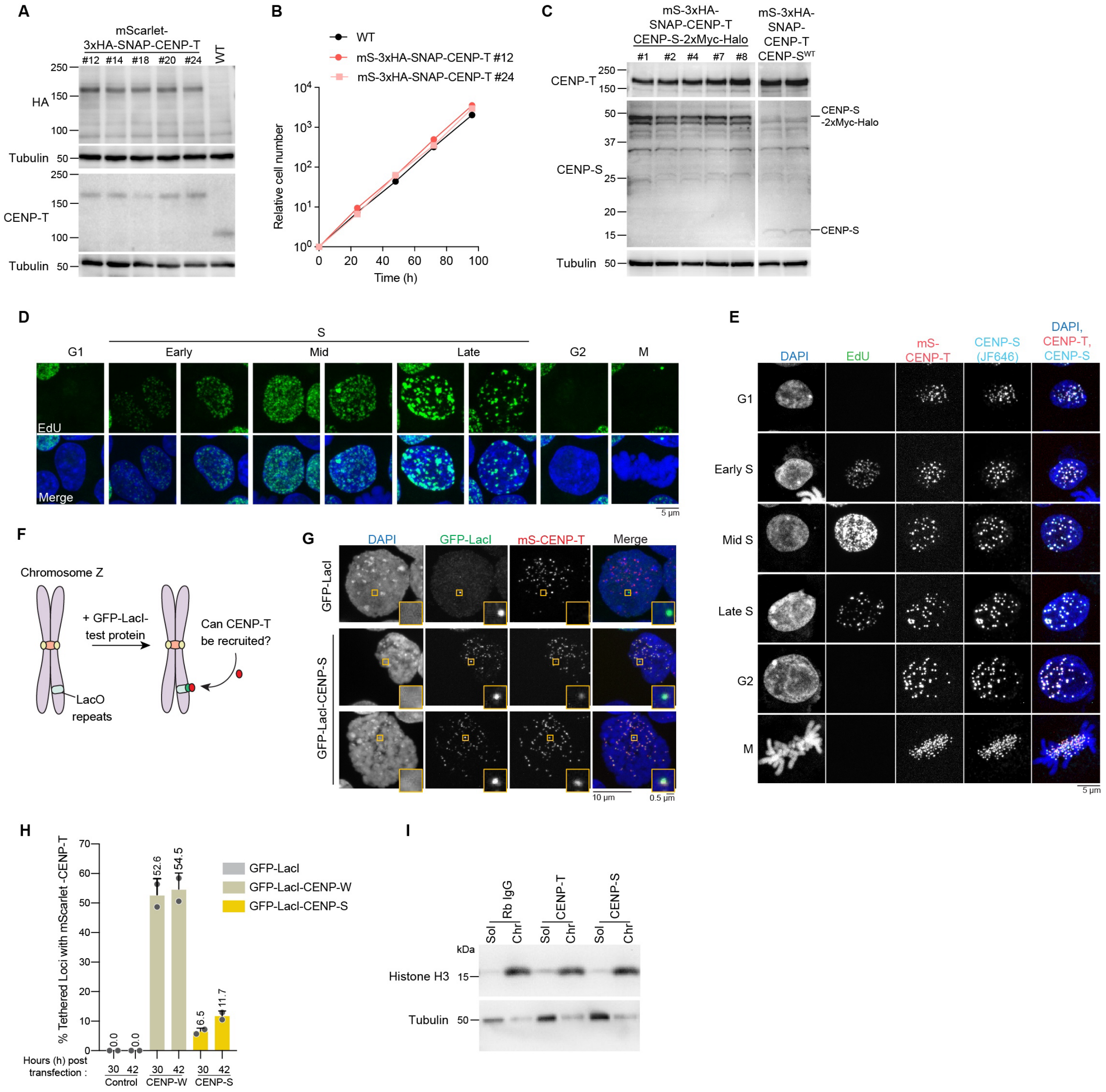

**Supplementary Figure 1. Validation of CENP-T complex formation and assembly in S and G2, Related to Figure 1.**

(A) mScarlet-3xHA-SNAP-CENP-T (mS-SNAP-CENP-T) expression in wild-type (WT, CL18) cells.

(B) Growth of WT and mS-SNAP-CENP-T cells. The cell numbers were normalized to those at 0 h for each cell line.

(C) CENP-S-2x-Myc-Halo (CENP-S-Halo) expression in mS-SNAP-CENP-T cells.

(D) Representative EdU staining pattern across cell cycle stages.

(E) Localization of CENP-T complex components, mS-SNAP-CENP-T and CENP-S-Halo labeled with JF646, across cell cycle stages.

(F) Schematic of lacO-LacI-based tethering assay. Plasmids expressing GFP-LacI-based fusion constructs were transfected into cells containing lacO repeats integrated at chromosome Z. Ability to recruit mScarlet-CENP-T was monitored at 30 and 42 h post-transfection.

(G and H) Localization of mS-CENP-T following transient expression of GFP-LacI fusion constructs into cells containing lacO repeats integrated at chromosome Z. Percentage of tethered foci recruiting mS-CENP-T was quantitated in (H). Mean is reported, $n = 2$, $\geq 250$ cells mean $\pm$ SD.

(I) Western blot showing efficiency of cell fractionation using Histone H3 and Tubulin levels, Related to Figure 1I.

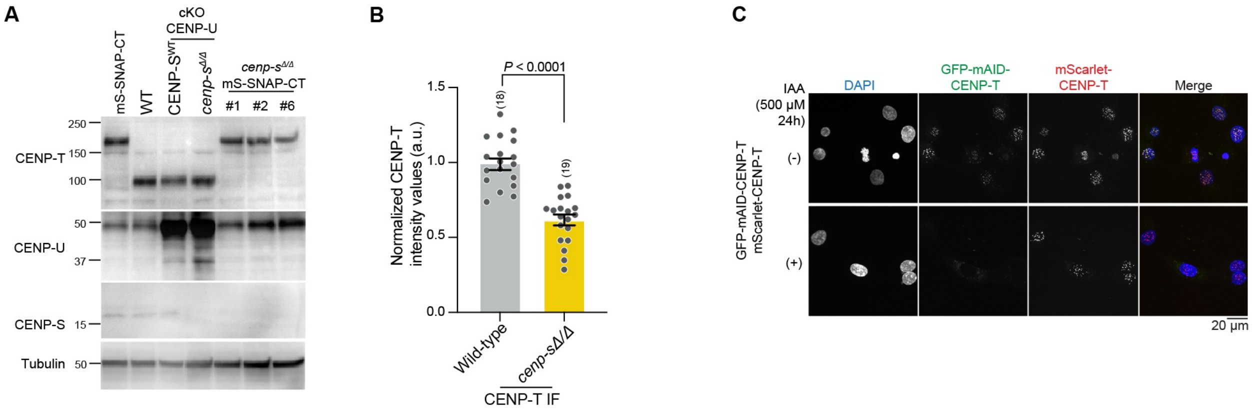


**Supplementary Figure 2. Strain validation and phenotypic confirmation supporting CENP-S-X role in CENP-T kinetochore stability, related to Figure 2**

(A) Presence of CENP-S protein in mS-SNAP-CENP-T, WT, cKO-CENP-U, cKO-CENP-U *cenp-s*$^{\Delta/\Delta}$ and mS-SNAP-CENP-T *cenp-s*$^{\Delta/\Delta}$ was examined.

(B) CENP-T levels in WT and *cenp-s*$^{\Delta/\Delta}$ was examined by immunofluorescence (IF) using anti-CENP-T antibody and quantified in mitotic cells. CENP-T signal was quantified and internally normalized to co-mixed reference cells. Statistical analysis was performed with a non-parametric t-test comparing two unpaired groups (Mann-Whitney test), number ($n$) of mitotic cells quantified is labelled in parenthesis, mean ± SEM, $P$ values are reported as indicated.

(C) Localization of CENP-T in human RPE-1 cells expressing GFP-mAID-CENP-T and mScarlet-CENP-T with and without IAA addition.

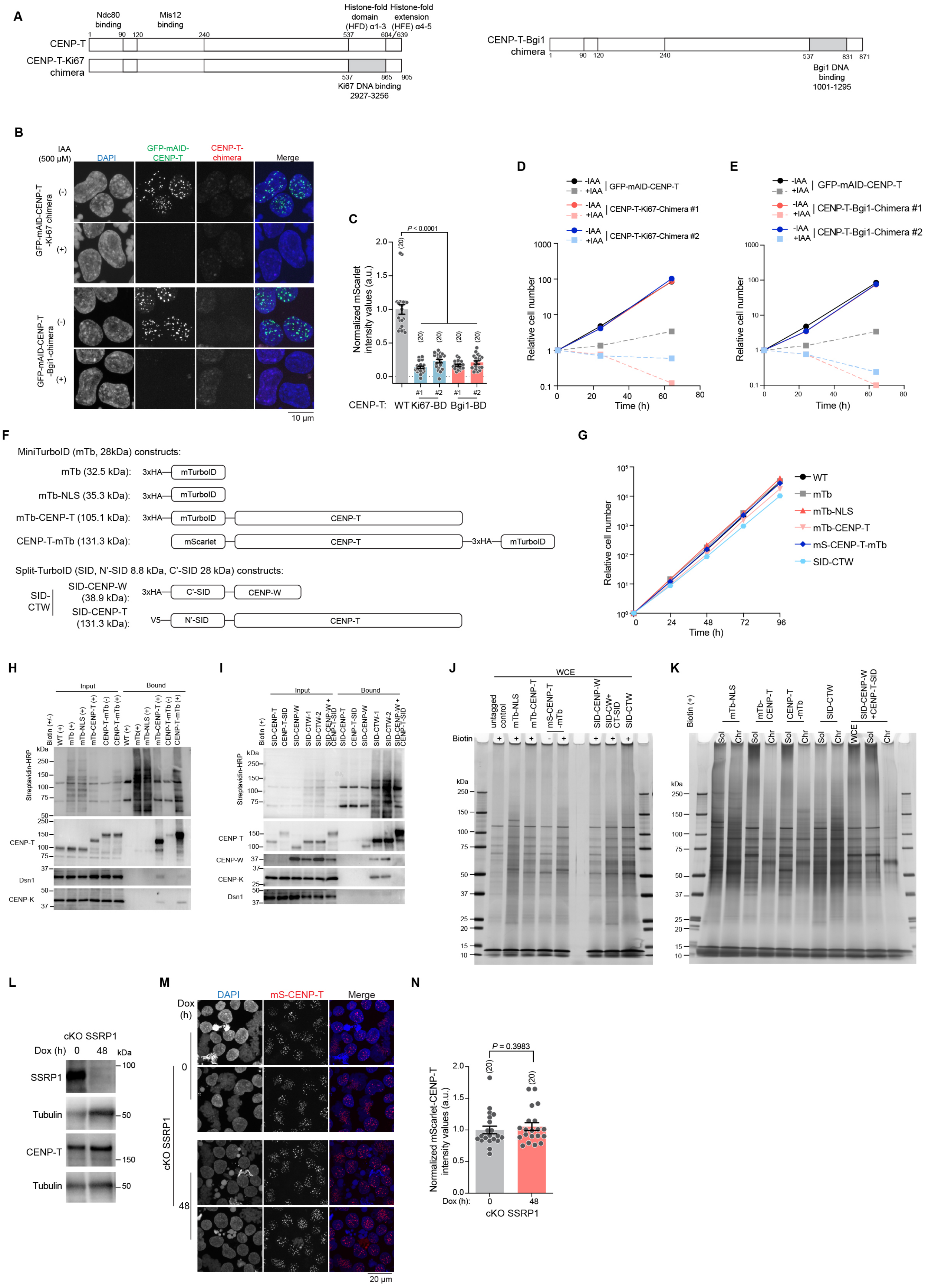

A
Ndc80 binding
Mis12 binding
Histone-fold domain (HFD) α1-3
Histone-fold extension (HFE) α4-5
1 90 120 240 537 604 639
CENP-T
CENP-T-Ki67 chimera
537 865 905
Ki67 DNA binding 2927-3256
CENP-T-Bgi1 chimera
1 90 120 240 537 831 871
Bgi1 DNA binding 1001-1295
B
IAA (500 μM)
DAPI
GFP-mAID-CENP-T
CENP-T-chimera
Merge
GFP-mAID-CENP-T -Ki67 chimera
GFP-mAID-CENP-T -Bgi1-chimera
(-)
(+)
10 μm
C
P < 0.0001
Normalized mScarlet intensity values (a.u.)
(20)
#1 #2 #1 #2
CENP-T: WT Ki67-BD Bgi1-BD
D
-IAA +IAA | GFP-mAID-CENP-T
-IAA +IAA | CENP-T-Ki67-Chimera #1
-IAA +IAA | CENP-T-Ki67-Chimera #2
Relative cell number
Time (h)
E
-IAA +IAA | GFP-mAID-CENP-T
-IAA +IAA | CENP-T-Bgi1-Chimera #1
-IAA +IAA | CENP-T-Bgi1-Chimera #2
Relative cell number
Time (h)
F
MiniTurboID (mTb, 28kDa) constructs:
mTb (32.5 kDa): 3xHA mTurboID
mTb-NLS (35.3 kDa): 3xHA mTurboID
mTb-CENP-T (105.1 kDa): 3xHA mTurboID CENP-T
CENP-T-mTb (131.3 kDa): mScarlet CENP-T 3xHA mTurboID
Split-TurboID (SID, N'-SID 8.8 kDa, C'-SID 28 kDa) constructs:
SID-CTW
SID-CENP-W (38.9 kDa): 3xHA C'-SID CENP-W
SID-CENP-T (131.3 kDa): V5 N'-SID CENP-T
G
WT
mTb
mTb-NLS
mTb-CENP-T
mS-CENP-T-mTb
SID-CTW
Relative cell number
Time (h)
H
Input
Bound
Biotin (+/-)
WT (+)
mTb (+)
mTb-NLS (+)
mTb-CENP-T (+)
CENP-T-mTb (-)
CENP-T-mTb (+)
Streptavidin-HRP
CENP-T
Dsn1
CENP-K
kDa
I
Input
Bound
Biotin (+)
SID-CENP-T
CENP-T-SID
SID-CENP-W
SID-CTW-1
SID-CTW-2
SID-CENP-W + CENP-T-SID
Streptavidin-HRP
CENP-T
CENP-W
CENP-K
Dsn1
kDa
J
WCE
untagged control
mTb-NLS
mTb-CENP-T
mS-CENP-T -mTb
SID-CENP-W
SID-CW+ CT-SID
SID-CTW
Biotin
kDa
K
Biotin (+)
mTb-NLS
mTb-CENP-T
CENP-T -mTb
SID-CTW
SID-CENP-W +CENP-T-SID
Sol
Chr
WCE
kDa
L
cKO SSRP1
Dox (h) 0 48
kDa
SSRP1
Tubulin
CENP-T
Tubulin
M
Dox (h)
DAPI
mS-CENP-T
Merge
0
48
cKO SSRP1
20 μm
N
P = 0.3983
Normalized mScarlet-CENP-T intensity values (a.u.)
(20)
Dox (h): 0 48
cKO SSRP1

**Supplementary Figure 3. Chimeric mutant insufficiency towards CENP-T kinetochore targeting and biotin proximity labeling methodology validation, Related to Figure 3.**

(A) Schematic representation of chimeric CENP-T mutants. $\alpha$-helix 1-3 of the CENP-T histone fold domain was replaced with the basic DNA binding motifs from either human Ki67 or *Cryptococcus neoformans* Bridgin protein.

(B and C) Localization of stably expressed chimeric CENP-T in cells expressing GFP-mAID-CENP-T in the presence or absence of IAA for 3 h. Kinetochore localization of chimeric CENP-T was measured at mitotic kinetochores in (C). Statistical analysis was performed with Kruskal–Wallis one-way analysis followed by Dunn's multiple comparison test, number (*n*) of mitotic cells quantified is labelled in parenthesis, mean $\pm$ SEM, *P* values are reported as indicated.

(D and E) Growth of parental (GFP-mAID-CENP-T) cells and CENP-T-Ki67 chimeras in (D) and CENP-T-Bgi1 chimeras in (E) with or without IAA . The cell numbers were normalized to those at 0 h for each cell line.

(F) Schematic representation of miniTurboID (mTb) constructs, mTb, mTb-NLS, mTb-CENP-T, CENP-T-mTb and split-TurboID (SID) constructs of CENP-T and CENP-W.

(G) Growth of WT (CL18), mTb, mTb-NLS, mTb-CENP-T, CENP-T-mTb and SID-CENP-TW cells. The cell numbers were normalized to those at 0 h for each cell line.

(H) Western blot analysis of proteins biotinylated in mTb expressed cells. WT untagged control, mTb, mTb-NLS, mTb-CENP-T and CENP-T-mTb expressing cells were treated with biotin (50 $\mu$M) for 6 h or without and biotin affinity pull-down (AP) was performed from whole cell extract (WCE).

(I) Western blot analysis of proteins biotinylated in SID expressed cells. SID-CENP-T, CENP-T-SID, SID-CENP-W, SID-CTW, and SID-CENP-W+CENP-T-SID expressing cells were treated with biotin (50 $\mu$M) for 24 h and biotin affinity pull-down (AP) was performed from whole cell extract (WCE).

(J and K) Silver stain visualization of biotinylated proteins isolated following AP from whole cell extract (J), soluble and chromatin fractions (K) from mTb and SID expressing cells.

(L) Expression of SSRP1 in Tet-Off SSRP1 (cKO SSRP1) cells grown with or without doxycycline (Dox).

(M and N) Localization of CENP-T in cKO SSRP1 grown with or without Dox. mScarlet-CENP-T levels were quantified in mitotic cells. Statistical analysis was performed with a non-parametric t-test comparing two unpaired groups (Mann-Whitney test), number (*n*) of mitotic cells quantified is labelled in parenthesis, mean $\pm$ SEM, *P* values are reported as indicated.

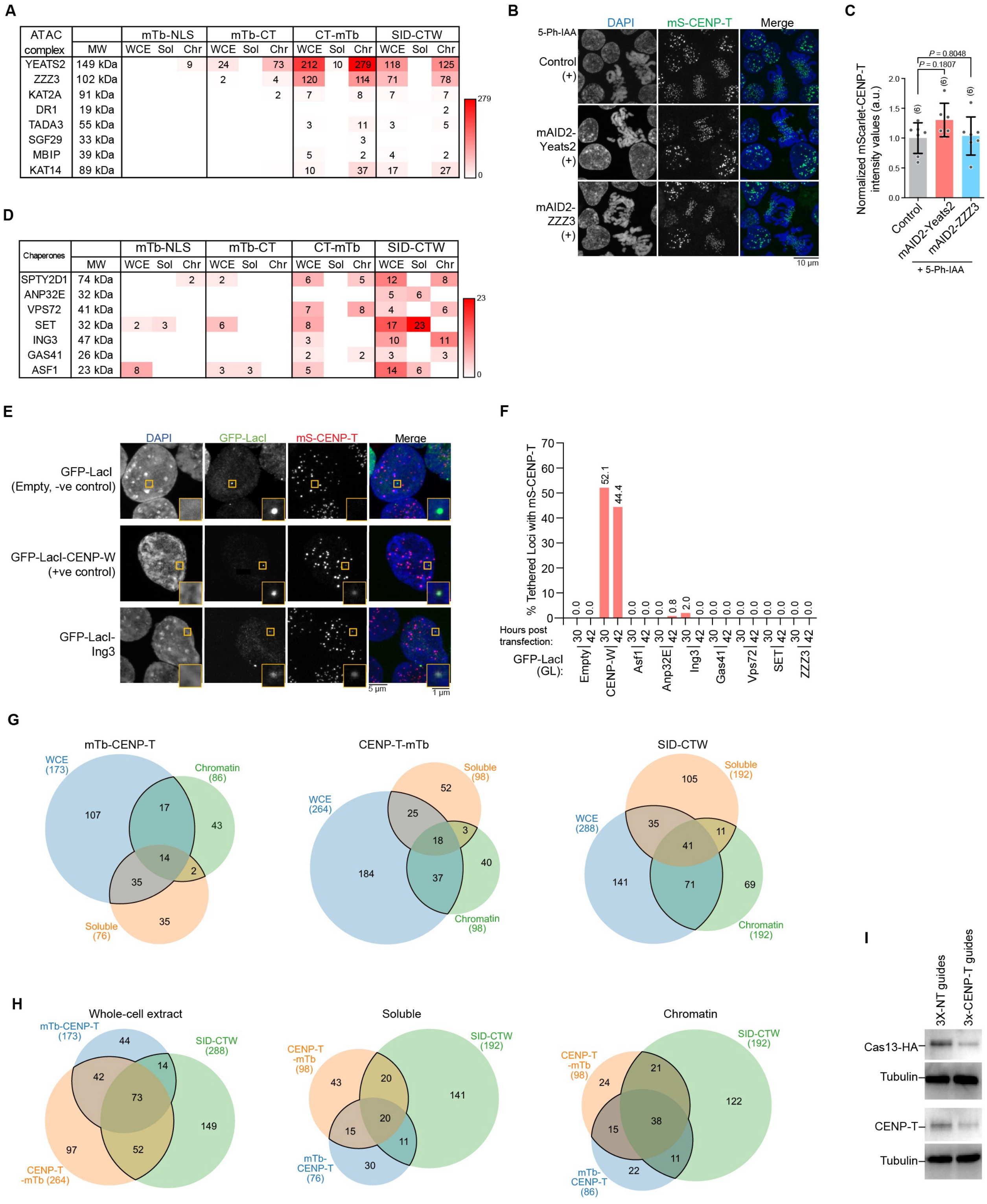


| ATAC complex | MW | mTb-NLS WCE | mTb-NLS Sol | mTb-NLS Chr | mTb-CT WCE | mTb-CT Sol | mTb-CT Chr | CT-mTb WCE | CT-mTb Sol | CT-mTb Chr | SID-CTW WCE | SID-CTW Sol | SID-CTW Chr |
|---|---|---|---|---|---|---|---|---|---|---|---|---|---|
| YEATS2 | 149 kDa | | | 9 | 24 | | 73 | 212 | 10 | 279 | 118 | | 125 |
| ZZZ3 | 102 kDa | | | | 2 | | 4 | 120 | | 114 | 71 | | 78 |
| KAT2A | 91 kDa | | | | | | 2 | 7 | | 8 | 7 | | 7 |
| DR1 | 19 kDa | | | | | | | | | | | | 2 |
| TADA3 | 55 kDa | | | | | | | 3 | | 11 | 3 | | 5 |
| SGF29 | 33 kDa | | | | | | | | | 3 | | | |
| MBIP | 39 kDa | | | | | | | 5 | | 2 | 4 | | 2 |
| KAT14 | 89 kDa | | | | | | | 10 | | 37 | 17 | | 27 |

| Chaperones | MW | mTb-NLS WCE | mTb-NLS Sol | mTb-NLS Chr | mTb-CT WCE | mTb-CT Sol | mTb-CT Chr | CT-mTb WCE | CT-mTb Sol | CT-mTb Chr | SID-CTW WCE | SID-CTW Sol | SID-CTW Chr |
|---|---|---|---|---|---|---|---|---|---|---|---|---|---|
| SPTY2D1 | 74 kDa | | | 2 | 2 | | | 6 | | 5 | 12 | | 8 |
| ANP32E | 32 kDa | | | | | | | | | | 5 | 6 | |
| VPS72 | 41 kDa | | | | | | | 7 | | 8 | 4 | | 6 |
| SET | 32 kDa | 2 | 3 | | 6 | | | 8 | | | 17 | 23 | |
| ING3 | 47 kDa | | | | | | | 3 | | | 10 | | 11 |
| GAS41 | 26 kDa | | | | | | | 2 | | 2 | 3 | | 3 |
| ASF1 | 23 kDa | 8 | | | 3 | 3 | | 5 | | | 14 | 6 | |

**Supplementary Figure 4. Biotin proximity screen validation and characterization of CENP-T-W interaction hits, Related to Figure 3.**

(A) Total spectral counts for ATAC complex components detected in the biotin proximity labeling screens described in Figure 3D.

(B and C) Kinetochore localization of CENP-T in mScarlet-mAID2-Yeats2 or mScarlet-mAID2-ZZZ3 with 5-Ph-IAA for 8 h. mScarlet CENP-T signal in mitotic cells was quantified in (C). Statistical analysis was performed with Kruskal–Wallis one-way analysis followed by Dunn's multiple comparison test, number (*n*) of mitotic cells quantified is labelled in parenthesis, mean ± SEM, *P* values are reported as indicated.

(D) Total spectral counts for histone chaperons detected in the biotin proximity labeling screens described in Figure 3D.

(E and F) Localization of mS-CENP-T following transient expression of GFP-LacI (GL) fusion constructs of identified chaperone proteins from the biotin proximity labeling screen. Percentage of tethered foci recruiting mS-CENP-T was quantitated in (F). ≥ 100 cells.

(G and H) Number of hits identified by mass spectrometry (MS) in the WCE, soluble, or chromatin fractions of a single cell line (G), or across mTb-CENP-T, CENP-T-mTb, and SID-CTW cells for a given fraction (H) following biotin affinity purification. These hits were further screened as described in Figure 3G prior to the Cas13-based secondary microscopic screen.

(I) CENP-T levels in Tet-On Cas13 cells following the transfection of Cas13-GFP plasmid containing 3 pre-guides against non-target (NT) control or CENP-T gene.

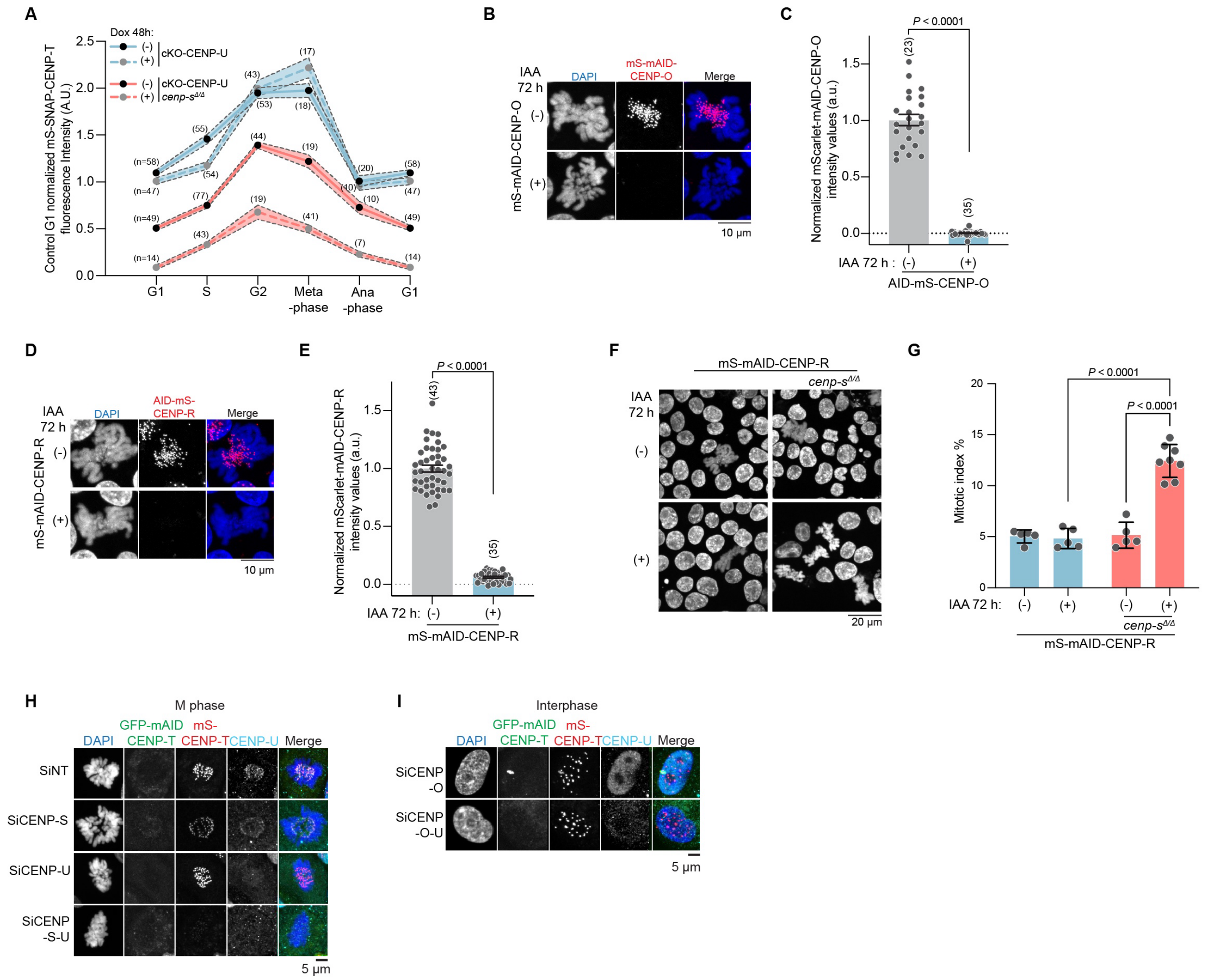

**Supplementary Figure 5. CENP-O complex mutant strain validation and cross-species conservation of CENP-S-X and CENP-O complex mediated CENP-T regulation, Related to Figure 4.**

(A) mS-SNAP-CENP-T signal intensities of cKO-CENP-U and cKO-CENP-U *cenp-s*$^{\Delta/\Delta}$, with and without Dox was quantified across cell cycle stages. Number (*n*) of cells quantified is labelled in parenthesis, mean ± SEM.

(B and C) Localization mS-mAID-CENP-O with and without IAA. CENP-O levels were quantified in mitotic cells. Statistical analysis was performed with a non-parametric t-test comparing two unpaired groups (Mann-Whitney test), number (*n*) of mitotic cells quantified is labelled in parenthesis, mean ± SEM, *P* values are reported as indicated.

(D and E) Localization mS-mAID-CENP-R with and without IAA. CENP-R levels were quantified in mitotic cells. Statistical analysis was performed with a non-parametric t-test comparing two unpaired groups (Mann-Whitney test), number (*n*) of mitotic cells quantified is labelled in parenthesis, mean ± SEM, *P* values are reported as indicated.

(F and G) Mitotic index of mS-mAID-CENP-R and mS-mAID-CENP-R *cenp-s*$^{\Delta/\Delta}$ with and without IAA, 72 h. Representative images of DAPI-stained nuclei are shown in (F). Mitotic index was quantified in (G). Statistical analysis was performed with Kruskal–Wallis one-way analysis followed by Dunn's multiple comparison test, mean ± SD, $n \geq 5$, > 1000 cells, *P* values are reported as indicated.

(H) Localization of CENP-T in mitotic human RPE-1 cells following treatment of SiRNA towards control (non-target guide, SiNT), CENP-S (SiCENP-S), CENP-U (siCENP-U) or both CENP-S and CENP-U (SiCENP-S-U). Related to Figure 4I.

(I) Localization of CENP-T in human RPE-1 cells following treatment of SiRNA towards CENP-O (siCENP-O) or both CENP-O and CENP-U (SiCENP-O-U). Related to Figure 4I.

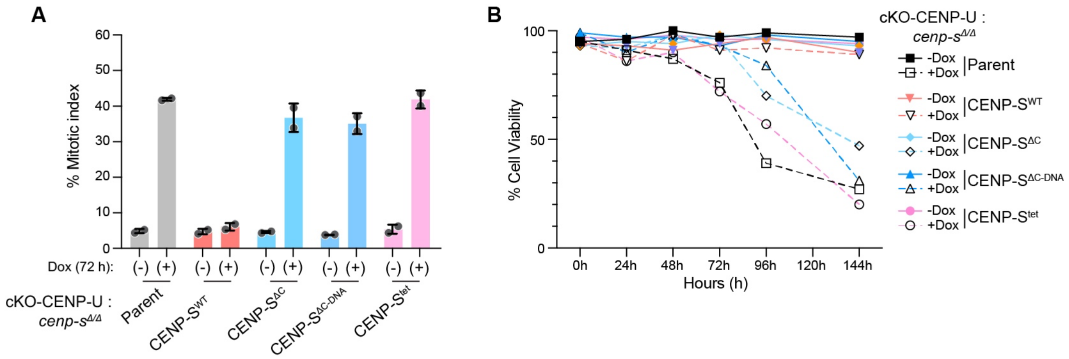


**Supplementary Figure 6. DNA-binding activity of CENP-S is essential in CENP-S/-U double mutants, Related to Figure 5.**

(A) Mitotic index of parental (cKO-CENP-U *cenp-s*$^{\Delta/\Delta}$) and the reintegration strains of CENP-S$^{FL}$, CENP-S$^{\Delta C}$, CENP-S$^{\Delta C+DNA}$ or CENP-S$^{tet}$, with and without Dox was quantified. Mean ± SD, *n* = 2, > 400 cells.

(B) Cell viability of parental (cKO-CENP-U *cenp-s*$^{\Delta/\Delta}$) and the reintegration strains of CENP-S$^{FL}$, CENP-S$^{\Delta C}$, CENP-S$^{\Delta C+DNA}$ or CENP-S$^{tet}$, with and without Dox was quantified.

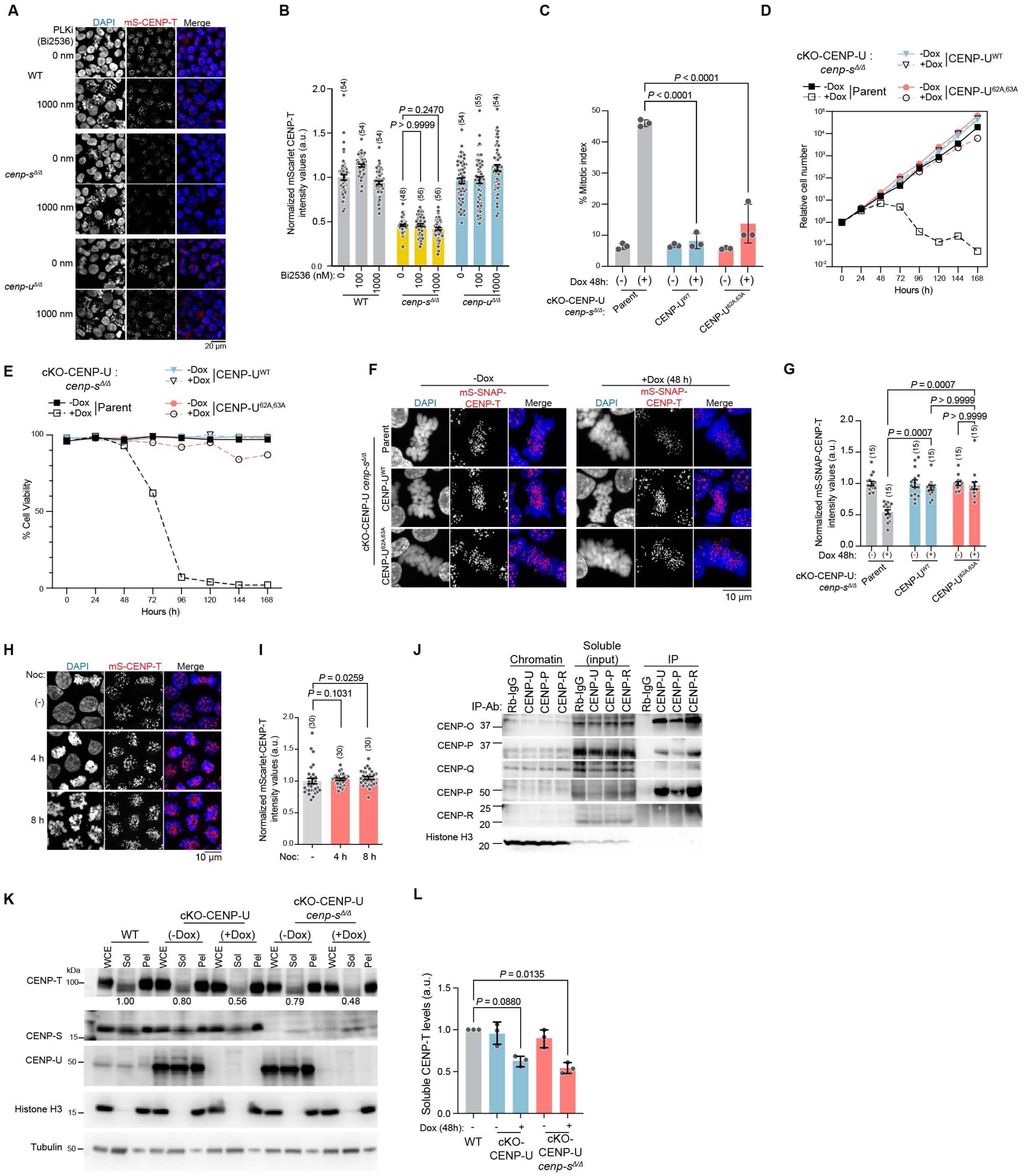


**Supplementary Figure 7. Canonical roles of the CENP-O complex components do not influence CENP-T complex stability, Related to Figure 6**

(A and B) Localization of mScarlet-CENP-T in WT (CL18), *cenp-s*$^{\Delta/\Delta}$ and *cenp-U*$^{\Delta/\Delta}$ cells following treatment with PLKi inhibitor Bi2536 at the indicated concentrations for 4 h. CENP-T signal in mitotic cells was quantified in (B). Statistical analysis was performed with Kruskal–

Wallis one-way analysis followed by Dunn's multiple comparison test, number (*n*) of mitotic cells quantified is labelled in parenthesis, mean ± SEM, *P* values are reported as indicated.

(C, D and E) Mitotic index (C), growth (D) and cell viability (E) of parental cKO-CENP-U *cenp-s*$^{\Delta/\Delta}$ cells and after reintegration of the CENP-U$^{WT}$ or CENP-U$^{62A,63A}$ construct was quantified with or without Dox treatment for 48 h. Statistical analysis was performed with Kruskal–Wallis one-way analysis followed by Dunn's multiple comparison test, *n* = 3, > 650 cells in (C). In (D) the cell numbers were normalized to those at 0 h for each cell line.

(F and G) Localization of mS-SNAP-CENP-T in parental cKO-CENP-U *cenp-s*$^{\Delta/\Delta}$ cells and after reintegration of the CENP-U$^{WT}$ or CENP-U$^{62A,63A}$ construct, with or without Dox treatment for 48 h. CENP-T signal in mitotic cells was quantified in (G). Statistical analysis was performed with Kruskal-Wallis one-way analysis followed by Dunn's multiple comparison test, number (*n*) of mitotic cells quantified is labelled in parenthesis, mean ± SEM, *P* values are reported as indicated.

(H and I) Localization of mS-CENP-T in WT (CL18) cells with or without Noc treatment (100 nM). CENP-T signal in mitotic cells was quantified in (G). Statistical analysis was performed with Kruskal-Wallis one-way analysis followed by Dunn's multiple comparison test, number (*n*) of mitotic cells quantified is labelled in parenthesis, mean ± SEM, *P* values are reported as indicated.

(J) Western blot of CENP-U, CENP-P and CENP-R immunoprecipitation (IP) from the soluble pool. Proteins were extracted following cell fractionation of WT (CL18) cells and IP was performed using anti-CENP-U, anti-CENP-P, anti-CENP-R or control Rabbit (Rb) IgG.

(K and L) CENP-T protein levels across whole-cell extract (WCE), soluble (Sol) and chromatin (Chr) pools in WT (CL18), cKO-CENP-U and cKO-CENP-U *cenp-s*$^{\Delta/\Delta}$ cells, with or without Dox treatment for 48 h. The signal intensity of CENP-T in the soluble pool was quantified in (H). Statistical analysis was performed with Kruskal-Wallis one-way analysis followed by Dunn's multiple comparison test, mean ± SD, *n* = 3, *P* values are reported as indicated.
